# Micro-C reveals MORC/ApiAP2-mediated links between distant, functionally related genes in the human malaria parasite

**DOI:** 10.1101/2024.08.28.610079

**Authors:** Parul Singh, Jacques Serizay, Justine Couble, Maureen D. Cabahug, Catarina Rosa, Patty Chen, Artur Scherf, Romain Koszul, Sebastian Baumgarten, Jessica M. Bryant

## Abstract

Genome organization plays a significant role in silencing heterochromatinized genes in the most virulent human malaria parasite, *Plasmodium falciparum*. However, it remains unclear how heterochromatinized genes spatially cluster or if active genes are also organized in a specific manner. We used Micro-C to achieve a near-nucleosome resolution DNA-DNA contact map, which revealed new inter- and intrachromosomal heterochromatic and euchromatic structures in the blood stage parasite. We observed subtelomeric fold structures that facilitate interactions amongst heterochromatinized genes involved in antigenic variation. In addition, we identified long-range intra- and interchromosomal interactions amongst active, stage-specific genes. Both structures are mediated by AP2-P, an ApiAP2 transcription factor, and a putative MORC chromatin remodeler, and functional specificity is achieved via combinatorial binding with other sequence-specific DNA-binding factors. This study provides unprecedented insight into the organizational machinery used by this medically important eukaryotic parasite to spatially coordinate genes underlying antigenic variation and to co-activate stage-specific genes.

## Introduction

Genome organization within the nucleus is important for transcriptional regulation and genome integrity of eukaryotes^1,2^. The simplest unit of eukaryotic genome organization is the nucleosome, which can impact gene transcription via positioning along the DNA or via the constituent histones and their post translational modifications. Eukaryotic genomes can present multiple organizational features including, but not limited to, telomere clustering, rDNA-containing nucleoli, active and inactive compartments, arrays of loops involved in the compaction of mitotic or meiotic chromosomes, or also extremely compacted genomes in certain male gametes^3^. A comprehensive body of work now points to the strong interplay between transcriptional activity and 3D organization of the genome^4^.

Contact maps of large animal genomes generated using the chromosome conformation capture (Hi-C^5^) approach display topologically associating domains, or TADs^6–9^. TADs are self-interacting regions in which loci tend to make more contacts with each other than with neighboring loci outside the region. In mammals, TADs appear as the reflection of the dynamic extrusion of large chromatin loops by the structural maintenance of chromosomes (SMC) cohesin complex^10,11^. CCCTC-Binding Factor (CTCF) molecules are major roadblocks to cohesin-mediated loop expansion in mammals, delimiting TAD boundaries bridged by a looping signal^12^. These loops are believed to promote and restrict interactions amongst genes and their cis-regulatory elements. However, CTCF/cohesin-independent enhancer-promoter contacts that require RNA polymerase II for their formation have also been described in mammals^13,14^. Transcription itself regulates the structure of the chromosome on a large scale, notably by interacting with loop extrusion^14–16^, but also on a short scale. Indeed, active transcription results in the formation of transcription-induced domains (TIDs) in bacterial^17^, yeast ^18^, and mammalian^19^ genomes, presumably through a local insulating effect of the RNA polymerase that favors DNA contacts between adjacent loci.

Chromatin spatial compartmentalization has been correlated with different transcriptional outcomes. Lamin-associated chromatin domains tend to encompass developmentally regulated or lowly expressed genes and have repressive chromatin signatures^20^. On the other hand, structures such as the nucleolus and nuclear speckles are located towards the interior of the nucleus and are associated with high levels of gene activity^21^. A large body of evidence has demonstrated that cohorts of active genes on the same and different chromosomes can physically associate in the nucleus and often overlap with RNA polymerase II-enriched foci, in what were called “transcription factories”^22–24^. Regardless of the term used, spatial association of genes in nuclear bodies is not random but mediated by specific chromatin-associated factors to achieve a boost in transcriptional activity^21^.

While most studies of genome organization have been performed in model organisms, non-model organisms such as protozoan pathogens offer fascinating insight into how nuclear architecture and transcription are connected to promote survival in a hostile environment. The life cycle of the most virulent human malaria parasite, *Plasmodium falciparum,* is driven by a complex transcriptional cascade in which each stage has a characteristic transcriptional program^25,26^. The influence of genome organization on transcription has been explored using Hi-C, which yielded genome-wide contact maps with 10-25 kb resolution^27,28^. These studies confirmed that the structure of this 23 Mb haploid genome is driven by the clustering of 1) rDNA loci^27–29^, 2) telomeric regions^27,28,30,31^, and 3) centromeres^27,28,32^, all features that were previously described in other eukaryotes and are important for genome stability, transcription, and compaction^33,34^.

However, this parasite exhibits a genome organization that underlies its pathogenicity and interaction with its human host. A remarkable feature of its genome architecture is the strong inter- and intrachromosomal association of heterochromatinized genes that are involved in antigenic variation, pathogenesis, and sexual development^27,35–37^. These genes include *ap2-g*, a transcription factor involved in the initiation of sexual commitment (gametocytogenesis) (**Fig. 1A**), and virulence genes (such as *var*) that belong to multigene families, which encode variant surface antigens that are crucial to infection and pathogenesis. Whether they are located in subtelomeric or central chromosomal regions, these genes are bound by heterochromatin protein 1 (HP1), and a subset forms clusters at the nuclear periphery^37,38^. This clustering is believed to be involved in the coordination of *var* gene mutually exclusive expression^27,35,39^ as well as recombination amongst *var* genes, which generates antigenic diversity^30,40–43^.

**Figure 1:**
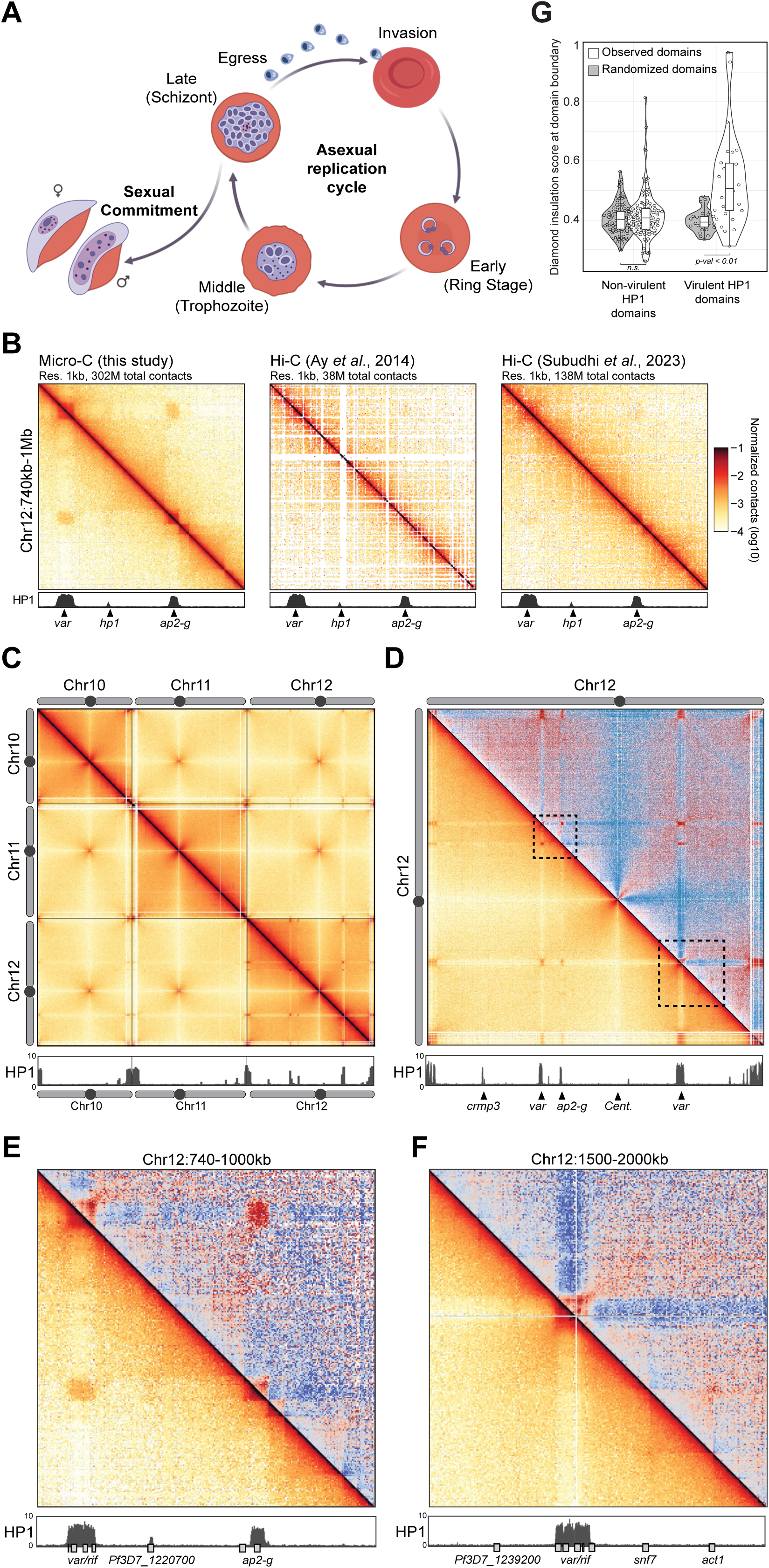
Micro-C provides a high-resolution view of *P. falciparum* genome organization. **A)** The asexual replication cycle begins when a merozoite that has just egressed from an infected human red blood cell invades a new red blood cell. During the cycle, the parasite develops from a ring (early stage) to a trophozoite (middle stage), which undergoes DNA replication and schizogony to form a schizont (late stage). Genes that are needed for egress of the merozoites from the infected red blood cell and invasion of a new red blood cell are expressed in late-stage parasites. A small percentage of parasites exit the asexual cycle to undergo sexual commitment through a process called gametocytogenesis, which generates male and female gametes that can be transmitted to the mosquito. Figure created with BioRender.com. **B)** Comparison of Micro-C data to previously published Hi-C datasets^27,66^ from *P. falciparum*. Contact map of a portion of Chromosome 12. HP1 ChIP/input ratio track from late-stage parasites^110^ is shown at the bottom with selected genes indicated below. **C-F)** Micro-C contact maps in late-stage parasites over (C) chromosomes 10 to 12 (5kb resolution), (D) the entire chromosome 12 (2kb resolution), (E) a 260kb-wide section of chromosome 12 (1kb resolution), and (F) a 500kb-wide section of chromosome 12 (2kb resolution). (E) and (F) are delineated with dashed line boxes in (D). (D-F) The bottom corner is normalized interaction frequency, and the top corner is log_2_-scaled observed/expected interaction frequency ratio. HP1 ChIP/input ratio track from late-stage parasites^110^ is shown at the bottom with selected genes indicated below. **G)** Insulation score at boundaries of HP1 central chromosomal domains, split according to whether each domain contains virulence genes. Randomly chosen domains were used as a control.

The transcriptional repression of these heterochromatinized genes is critical for maintaining an infection in the human host; however, other than HP1^37^, it is still unclear what protein factors underly their organization. Importantly, *P. falciparum* lacks CTCF^44^ and lamins^45^, which are important for genome organization in other eukaryotes. Moreover, the *P. falciparum* genome encodes relatively few sequence-specific DNA-binding factors, most of which belong to the 27-member ApiAP2 family. All ApiAP2 proteins contain at least one Apetala2/ERF DNA binding domain, which is also found in plant transcription factors^46^. ApiAP2 factors have been implicated in transcriptional regulation at multiple different stages of the parasite life cycle, but their role in genome organization has not been well established^47^.

The focus of most genome structure studies in *P. falciparum* has been on the organization of silent, heterochromatinized genes. However, Hi-C resolution remains limited to around 10 kb in the extremely AT-rich *P. falciparum* genome^48^, which is insufficient to identify finer-scale structure and leaves several questions open regarding: 1) the structuration of heterochromatin domains, 2) whether euchromatic genes form structured interactions like heterochromatic genes; 3) whether long-range chromatin loops form to bring distant loci (such as enhancers and promoters) in close proximity to contribute to gene regulation; and 4) which factors mediate these structures. To tackle these questions, we adapted Micro-C^18^, a nucleosome-based Hi-C derivative, to study *P. falciparum* genome organization at sub-kb resolution during the late stage of the asexual replication cycle, which takes place in the human red blood cells (**Fig. 1A**). The integration of these data with chromatin immunoprecipitation and sequencing (ChIP-seq) and RNA-sequencing (RNA-seq) data revealed that the *P. falciparum* genome is organized into heterochromatic and euchromatic domains whose boundaries are formed by ApiAP2 DNA-binding factors and a putative microorchidia chromatin remodeler, MORC. The variable composition of these complexes defines the function of different boundary elements: some facilitate folding of heterochromatic subtelomeric regions while others facilitate intra- and interchromosomal long-range interactions that create hubs of stage-specific transcription. This study reveals an unprecedented complexity of a small eukaryotic genome that relies on relatively few sequence-specific DNA-binding proteins to achieve drastic transcriptional changes that enable its complicated parasitic life cycle.

## Results

### Micro-C provides unprecedented resolution of *P. falciparum* genome organization

To generate a high-resolution genome-wide contact map in *P. falciparum* that allows for comparison to nucleosome-scale datasets (i.e. ChIP-seq and assay for transposase-accessible chromatin with sequencing (ATAC-seq)), we adapted Micro-C^18^ to this organism. Micro-C uses micrococcal nuclease (MNase) to fragment the genome down to the nucleosome level, providing a higher short-range resolution of chromatin conformation than the Hi-C protocol, which uses restriction enzymes. We performed four replicates in clonal wild-type parasites in the late stage of the red blood cell cycle, which showed high correlation and were subsequently combined to achieve higher resolution. We observed a strong nucleosome banding pattern in our Micro-C chimeric fragments (**Supp.** Fig. 1A), and the intra-chromosomal distance-dependent genomic interactions frequency P(s) profile (i.e., contact decay curve) revealed a typical polymer behavior with a linear P(s) (slope ∼-1.2) maintained for distances as short as 1 kb (**Supp.** Fig. 1B). Thus, we confirm that Micro-C substantially increased resolution of the *P. falciparum* contact map down to 1 kb (**Fig. 1B**).

We observed previously described interchromosomal interactions within and between telomeres and centromeres (**Supp.** Fig. 1C) and strong inter- and intrachromosomal interactions of virulence genes that are heterochromatinized by HP1, such as the ∼60-member *var* multigene family (**Fig. 1C**). In general, we did not observe strong inter- or intrachromosomal contacts between HP1-enriched virulence genes and HP1-enriched non-virulence genes such as *crmp3* (**Fig. 1D**). An exception is HP1-heterochromatinized *ap2-g*, shown to interact with HP1-enriched virulence genes (**Fig. 1D,E**)^37^. In addition, HP1 domains overlapping virulence genes (“virulent HP1 domains”, e.g., the central chromosomal *var* gene clusters on chromosomes 4, 7, 8, and 12 that encompass 4-7 *var* genes each), as well as an HP1 island comprised of a single gene, *ap2-g*, formed well-defined insulated domains (**Fig. 1D-G**). In contrast, HP1 domains containing other non-virulence genes (“non-virulent HP1 domains”, e.g., *crmp3*) do not form such insulated domains (**Fig. 1D,G**). Taken together, these data suggest that HP1 alone is not sufficient to insulate a heterochromatic domain from nearby euchromatin, nor to dictate the coalescence of distant genomic loci into a heterochromatin compartment.

### Micro-C reveals subtelomeric fold structures defined by a multiprotein complex

The high-resolution contact map allows the study of fine-scale organization within subtelomeric HP1-enriched domains in the *P. falciparum* genome. HP1 heterochromatin encompasses each subtelomeric region, including the non-genic region adjacent to the telomere as well as several virulence genes downstream that encode variant surface antigens (**Fig. 2A**; **Supp.** Fig. 2A). However, we detected a clear break in local contacts in the middle of this HP1 domain, forming two smaller self-associating domains: one non-genic and one genic (**Fig. 2A; Supp. Fig. 2B, left**). The non-genic HP1 subtelomeric domain starts just downstream of the telomere and ends just upstream of the first gene, which is usually a *var* gene (**Fig. 2A; Supp. Fig. 2A**). This domain forms a fold-like structure anchored at its boundaries, indicated by a corner point structure in the Micro-C data (**Fig. 2A,B; Supp. Fig. 2B, middle**). In contrast, genic HP1 subtelomeric domains do not form a fold-like structure (**Fig. 2A**). Importantly, we observe these structures only at chromosome ends that contain *var* genes, suggesting a link between this fold structure and *var* gene biology (**Supp.** Fig. 2C).

**Figure 2:**
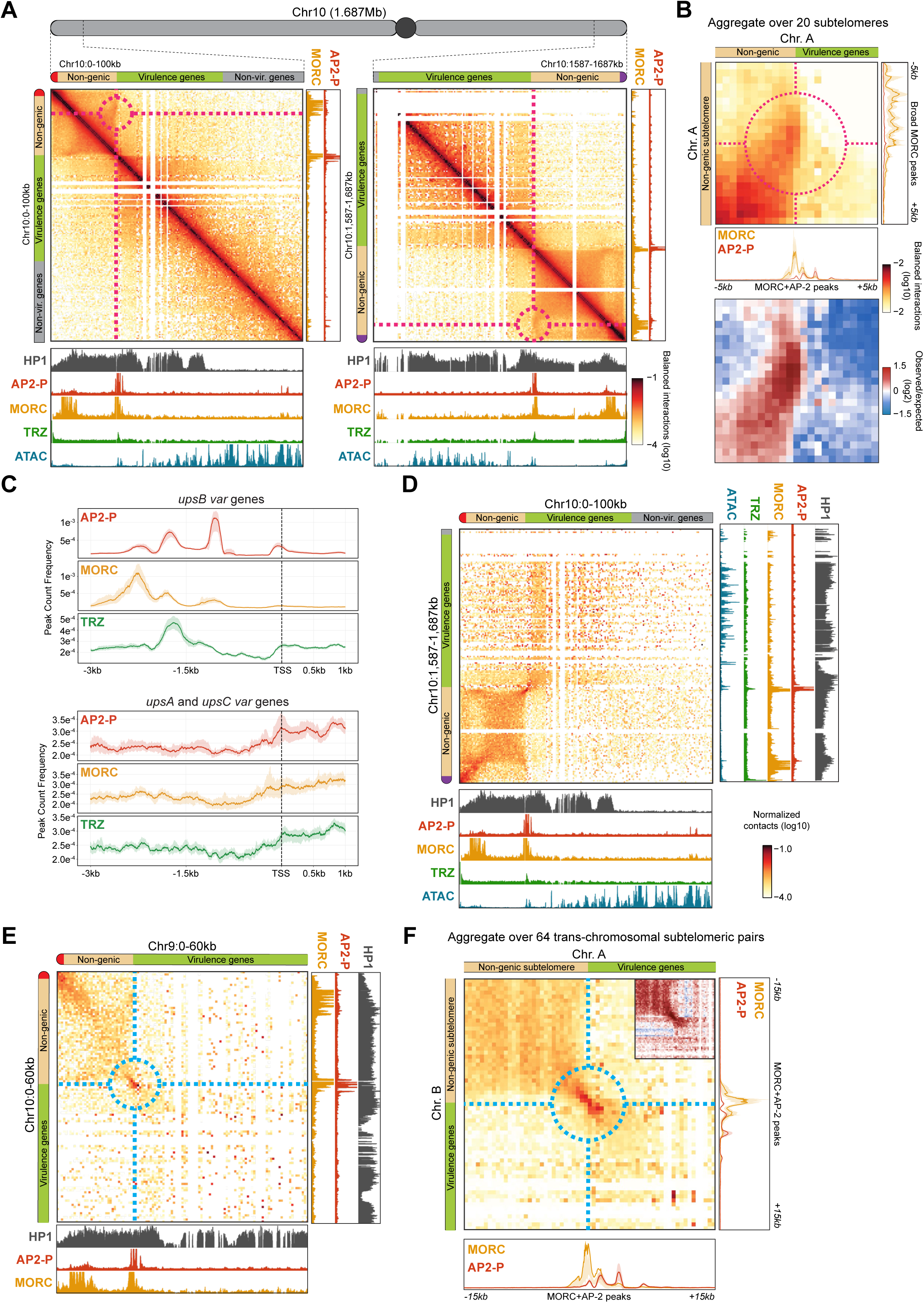
Micro-C reveals subtelomeric fold structures defined by a multiprotein complex. **A)** Micro-C contact maps of either end of Chromosome 10 in late-stage parasites (100kb-wide, 1kb resolution). AP2-P, MORC, HP1^110^, TRZ^60^ ChIP/input ratio tracks and ATAC-seq^58^ track from late-stage parasites are shown at the bottom and to the right of the contact maps. To the top and left of each contact map: red is the left arm telomere, tan is the non-genic subtelomeric region, green is the virulence gene-encoding subtelomeric region, grey is euchromatic genes, and purple is the right arm telomere. The dotted red circle indicates the subtelomeric fold contact point. **B)** Off-diagonal Micro-C contact maps aggregated over 20 subtelomeric loci (Table 10), characterized by the presence of a non-genic subtelomeric region (±5kb, 500bp resolution). For each subtelomeric locus, a 10kb-wide contact map was extracted, centered at the contact (indicated with red dotted circle) between the broad MORC peak in non-genic subtelomeres on one side, and MORC/AP2-P–enriched transition between non-genic and virulence gene–coding subtelomeric segment on the other side. This map highlights the fold-like structure detected by Micro-C. Contact maps extracted from subtelomeric loci located at the end of chromosomes have been mirrored so that the telomere is always positioned towards the top left corner. The top panel is normalized interaction frequency, and the bottom panel is log_2_-scaled observed/expected interaction frequency ratio. 1D aggregated AP2-P and MORC ChIP/input signals from late-stage parasites are shown below and on the right of the 2D aggregated Micro-C map. **C)** Metagene plots showing average AP2-P and MORC ChIP enrichment in clonal AP2-P-3HA-*glmS* and MORC-3HA parasites, respectively, in late-stage parasites from 3 kb upstream to 1 kb downstream of the predicted TSS for subtelomeric *var* genes that are transcribed in the opposite direction of the telomere (*upsB*, top) and subtelomeric *var* genes that are transcribed in the direction of the telomere (*upsA*) or central *var* genes (*upsC*, bottom). TRZ ChIP-seq data^60^ from stage-matched parasites are included. One replicate was used for the AP2-P and MORC ChIP datasets. **D)** Micro-C contact map between both subtelomeric regions of chromosome 10 in late-stage parasites (100kb-wide contact map, 1kb resolution), presented otherwise as in (Fig. 2A). **E)** Interchromosomal Micro-C contact map between subtelomeric regions of the left arm of chromosome 10 and of the left arm of chromosome 9 in late-stage parasites (60kb-wide, 1kb resolution), presented otherwise as in (Fig. 2A). The dotted blue circle indicates the subtelomeric contact point. **F)** Interchromosomal Micro-C contact maps aggregated over the 64 interchromosomal pairs of subtelomeric loci, characterized by the presence of a non-genic subtelomeric region (±15kb, 500bp resolution). For each interchromosomal pair of subtelomeric loci, a 30kb-wide contact map was extracted, centered at the MORC/AP2-P–enriched transition between non-genic and virulence gene-coding subtelomeric segment of each chromosome. Contact maps extracted from subtelomeric loci located at the end of chromosomes have been mirrored so that the telomere is always positioned towards the top left corner. Inset is log_2_-scaled observed/expected interaction frequency ratio of the area highlighted with a blue circle. 1D aggregated AP2-P and MORC ChIP/input signals from late-stage parasites are shown below and on the right of the 2D aggregated Micro-C map.

We next sought to identify the molecular factors structuring this subtelomeric fold. In other eukaryotes, SMC protein complexes facilitate the interaction of domain boundaries in the genome; however, SMC3 ChIP-seq data do not suggest that the cohesin complex plays this role in *P. falciparum*^49^. Incidentally, we recently identified proteins that are enriched in the chromatin of *var* gene promoters using a CRISPR-based proteomics approach^50^. Included in this protein cohort were two ApiAP2 DNA-binding factors – SPE2-interacting protein (SIP2, PF3D7_0604100) and AP2-P (PF3D7_1107800) – and a putative microorchidia (MORC, PF3D7_1468100) chromatin remodeler. In other eukaryotes, MORC proteins are ATPases that bind non-specifically to and condense DNA^51^. Thus, most MORC proteins, including in *Toxoplasma gondii*, have been implicated in gene repression and heterochromatin formation^52–57^.

We determined genome-wide binding of AP2-P and MORC by first generating epitope-tagged strains (**Supp.** Fig. 2D,E) and then performing ChIP-seq in clonal parasites during the late stage of red blood cell development (**Fig. 1A**), which also happens to be when these two proteins show highest expression (**Supp.** Fig. 2F,G). The ChIP-seq data confirmed the findings from our previous study^50^ and showed that the highest enrichment of AP2-P and MORC is upstream of subtelomeric *var* genes whose promoters are proximal to the telomere (known as *upsB var* genes, see **Supp.** Fig. 2A) (**Fig. 2A,B**; **Supp.** Fig. 2H; **Table 1**). We observe that MORC binds a broad region ∼8kb downstream of the telomere (mean 8.3kb ± 1.7kb), and jointly with AP2-P in three large, discreet peaks 1-2 kb upstream of the transcription start site (TSS) of almost all *upsB* subtelomeric *var* genes (**Fig. 2B,C**; **Supp.** Fig. 2H; **Table 1**). While all *var* genes are transcriptionally silent in late-stage parasites, we did not observe this same enrichment pattern upstream of subtelomeric *var* genes whose promoters are distal to the telomere (*upsA*) or central chromosomal *var* genes (*upsC*) (**Fig. 2C**; **Supp.** Fig. 2A). Importantly, we did not observe significant AP2-P and MORC enrichment at the boundaries between HP1 heterochromatin and euchromatin (**Fig. 2A**).

Despite being highly enriched in HP1, subtelomeric AP2-P/MORC-bound loci show significant chromatin accessibility, as measured by ATAC-seq^58^ (**Fig. 2A**; **Supp.** Fig. 2H). In fact, these same loci are also bound by other factors such as SIP2^50,59^ and telomere repeat-binding zinc finger protein (TRZ)^60^, which was shown to play a role in telomere homeostasis (**Fig. 2A,C**; **Supp.** Fig. 2H). Strikingly, the two regions of AP2-P and MORC subtelomeric enrichment precisely overlap the boundaries of the subtelomeric fold structure we identified with Micro-C (**Fig. 2A,B**). Enrichment of multiple factors at these subtelomeric structures where the genic region of the chromosome transitions to the non-genic subtelomere hints at a protein complex that could play an important role in the maintenance of chromatin structure.

We further found that subtelomeric AP2-P/MORC-bound loci form long-distance intrachromosomal interactions, identified by a focal point of contact between pairs of subtelomeric ends of individual chromosomes (**Fig. 2D**). These subtelomeric AP2-P/MORC-bound loci are also involved in trans-chromosomal contacts of subtelomeres (**Fig. 2E,F**; **Supp.** Fig. 2B, right; **Supp.** Fig. 2I). While HP1 is enriched in subtelomeric regions of chromosome 14, which lack *var* genes, AP2-P and MORC are not enriched there (**Supp.** Fig. 2C), and the above-mentioned intra- and interchromosomal subtelomeric contact points are not present (**Supp.** Fig. 2C,J). These data suggest that AP2-P and MORC facilitate intra- and interchromosomal interactions of subtelomeric regions that contain *var* genes.

### AP2-P is required for non-coding subtelomeric fold structures and interchromosomal interactions

To determine if AP2-P plays a role in *var* gene regulation or subtelomeric DNA structure, we performed a knockdown (KD) of this protein using an inducible *glmS* ribozyme system^61^. Despite a significant KD at the protein level (**Supp.** Fig. 3A), we did not observe an apparent growth phenotype or cell cycle arrest (**Supp.** Fig. 3B). Interestingly, AP2-P KD followed by RNA-seq and differential expression analysis in late-stage parasites (**Supp.** Fig. 3C; **Table 2**) confirmed that AP2-P KD did not affect cell cycle progression (**Supp.** Fig. 3D) and showed that there was no significant de-repression of *var* genes (**Supp.** Fig. 3E). However, AP2-P KD did result in the significant decrease of *morc* transcript levels (**Supp.** Fig. 3F). To determine if AP2-P KD results in MORC downregulation at the protein level, we added an epitope tag to *morc* in the AP2-P-3HA-*glmS* strain (**Supp.** Fig. 3G). In the resultant strain, AP2-P KD resulted in MORC downregulation in late-stage parasites (**Fig. 3A**). These data suggest that AP2-P regulates MORC and that an AP2-P KD is a MORC KD as well.

**Figure 3:**
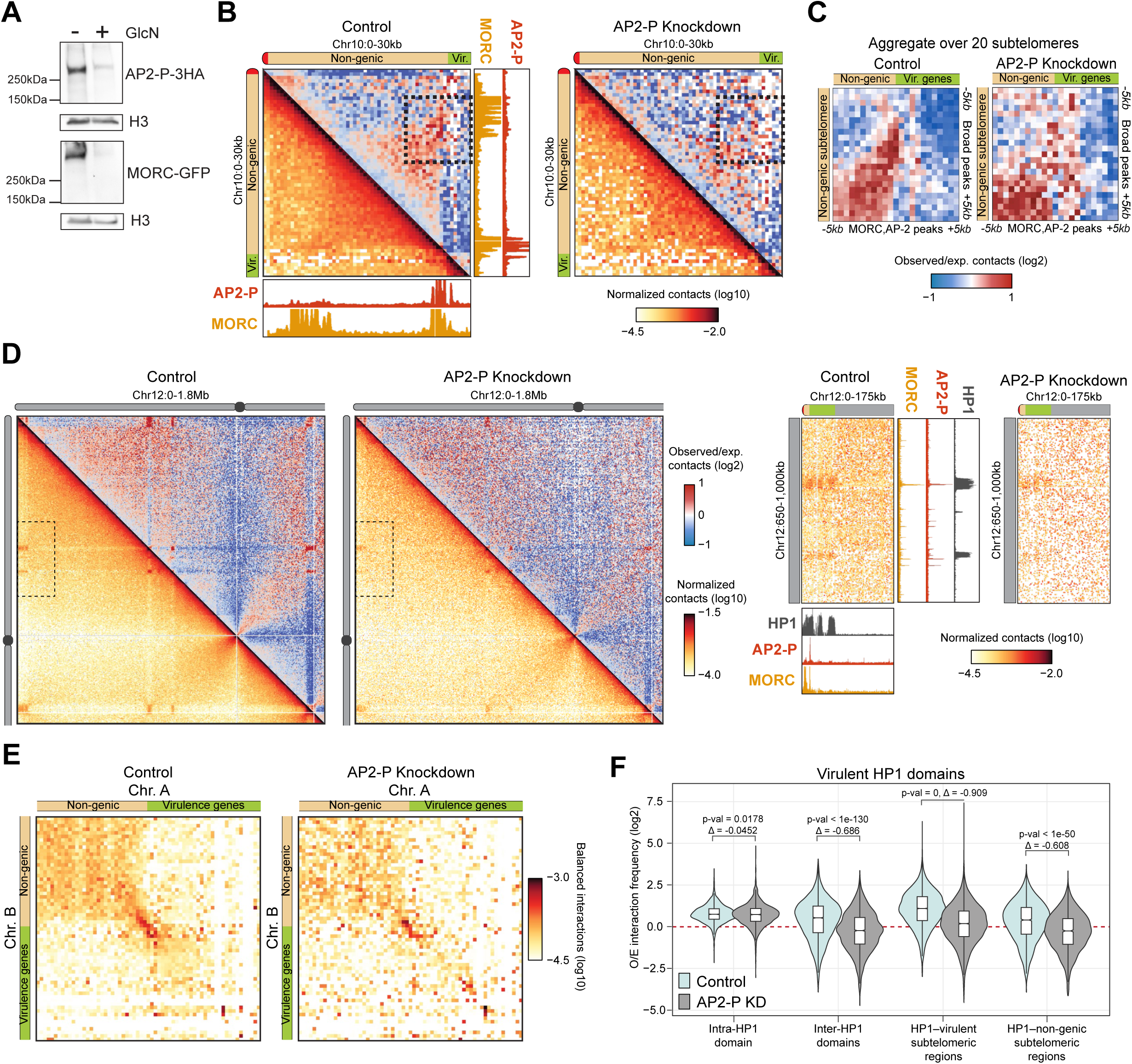
AP2-P is required for non-coding subtelomeric fold structures and interchromosomal interactions. **(A)** Western blot analysis of nuclear extracts from a synchronous clonal population of AP2-P-3HA-*glmS*:MORC-GFP late-stage parasites in the absence (-) or presence (+) of glucosamine (GlcN). Parasites were treated with glucosamine for two full cycles (96 h). AP2-P-3HA was detected with an anti-HA antibody, MORC-GFP was detected with an anti-GFP antibody, and an antibody against histone H3 was used as a loading control. Molecular weights are shown to the left. **(B)** For subsampled (65 million contacts) control (left) and AP2-P KD (right) Micro-C maps: Micro-C contact maps centered at the interaction of the left end of chromosome 10 in late-stage parasites (30kb-wide, 1kb resolution, bottom corner: normalized interaction frequency; top corner: log_2_-scaled observed/expected interaction frequency ratio), presented otherwise as in (Fig. 2A). The dotted black box indicates the area of contact between the subtelomeric fold anchor points. **(C)** For subsampled (65 million contacts) control (left) and AP2-P KD (right) Micro-C maps: off-diagonal Micro-C contact maps aggregated over 20 subtelomeric loci (Table 10) characterized by the presence of a non-genic subtelomeric region (±5kb, 500bp resolution), presented otherwise as in (Fig. 2B). The color scale indicates the log_2_-scaled observed/expected interaction frequency ratio. **(D)** Left: For subsampled (65 million contacts) control (left) and AP2-P KD (right) Micro-C maps: Micro-C contact maps over a 1.8Mb-wide section of chromosome 12 in late-stage parasites (5kb resolution, bottom corner: normalized interaction frequency; top corner: log_2_-scaled observed/expected interaction frequency ratio). Right: Zoomed Micro-C off-diagonal contact map in control and AP2-P KD, focusing on the contact of the subtelomeric region of the left arm of chromosome 12 with a central region of the chromosome containing virulence genes (area shown with dotted rectangle at left). Presented otherwise as in (Fig. 2A). **(E)** Interchromosomal Micro-C contact maps for subsampled (65 million contacts) control (left) or AP2-P KD (right) aggregated over the 64 interchromosomal pairs of subtelomeric loci, characterized by the presence of a non-genic subtelomeric region (±15kb, 500bp resolution). For each interchromosomal pair of subtelomeric loci, a 30kb-wide contact map was extracted, centered at the MORC/AP2-P–enriched transition between non-genic and virulence gene–coding subtelomeric segment of each chromosome. Contact maps extracted from subtelomeric loci located at the end of chromosomes have been mirrored so that the telomere is always positioned towards the top left corner. **(F)** Log_2_-scaled observed/expected interaction frequency ratio for (i) intrachromosomal interactions between two virulence gene-containing HP1 domains, (ii) interchromosomal interactions between two virulence gene-containing HP1 domains, (iii) interactions between virulence gene-containing HP1 domains and virulence gene-coding subtelomeric loci, or (iv) interactions between virulence gene-containing HP1 domains and non-genic subtelomeric loci. Contacts in the subsampled control dataset are shown in cyan and those in the AP2-P KD dataset are shown in grey.

While AP2-P (and thus MORC) downregulation did not substantially affect *var* gene transcription, AP2-P KD followed by Micro-C revealed significant structural consequences. First, AP2-P KD led to a loss of the subtelomeric fold structure upstream of subtelomeric *var* genes (**Fig. 3B**). In particular, the subtelomeric focal contacts between genomic loci bound by AP2-P and MORC were disrupted, suggesting that AP2-P is responsible for anchoring these subtelomeric fold structures (**Fig. 3C**). In addition, AP2-P KD resulted in a significant reduction in intra-(**Fig. 3D**) and interchromosomal (**Fig. 3E**) interactions of HP1-enriched regions containing virulence genes (such as the *var* gene family), but not other HP1-enriched regions (**Fig. 3F**; **Supp.** Fig. 3H). The reduction in trans-chromosomal contacts between subtelomeric regions was modest, suggesting that SIP2 or TRZ may also help to facilitate these interactions (**Fig. 3E**). Consistent with the fact that AP2-P and MORC are generally not enriched at the boundaries between euchromatin and HP1 heterochromatin (**Fig. 2A**), AP2-P KD did not affect insulation of HP1 self-associating domains that encompass virulence genes (**Supp.** Fig. 3I). Taken together, these data suggest that AP2-P and MORC play a role in the formation of long-range intra- and interchromosomal interactions between heterochromatic regions.

### AP2-P, AP2-I, and MORC activate stage-specific genes

In addition to their enrichment in subtelomeric heterochromatin, AP2-P and MORC show significant binding to sites within euchromatic regions of the genome that overlap with accessible chromatin (**Fig. 4A**; **Table 1**). Motif enrichment analysis of AP2-P ChIP-seq peaks identified a motif – “GTGCA” – that is shared with another ApiAP2 factor, AP2-I, which binds to the promoters of genes involved in late-stage parasite biology, such as red blood cell invasion^62^ (**Supp.** Fig. 4A). Comparison of AP2-I^62^ with AP2-P and MORC ChIP-seq data revealed significant overlap of these proteins throughout the genome (**Table 1**), with overlapping enrichment near the putative TSS of genes bound by AP2-P (**Fig. 4A,B**). Approximately half of all genes with an upstream peak of AP2-P also have an upstream peak of AP2-I, while 75% of all genes with an upstream peak of AP2-I also have an upstream peak of AP2-P (**Fig. 4C**; **Table 1**). Importantly, 90% of genes with an upstream peak of MORC also have an upstream peak of AP2-P, AP2-I, or both, suggesting a dependency of MORC binding on an ApiAP2 factor (**Fig.** 4C; Table 1).

**Figure 4:**
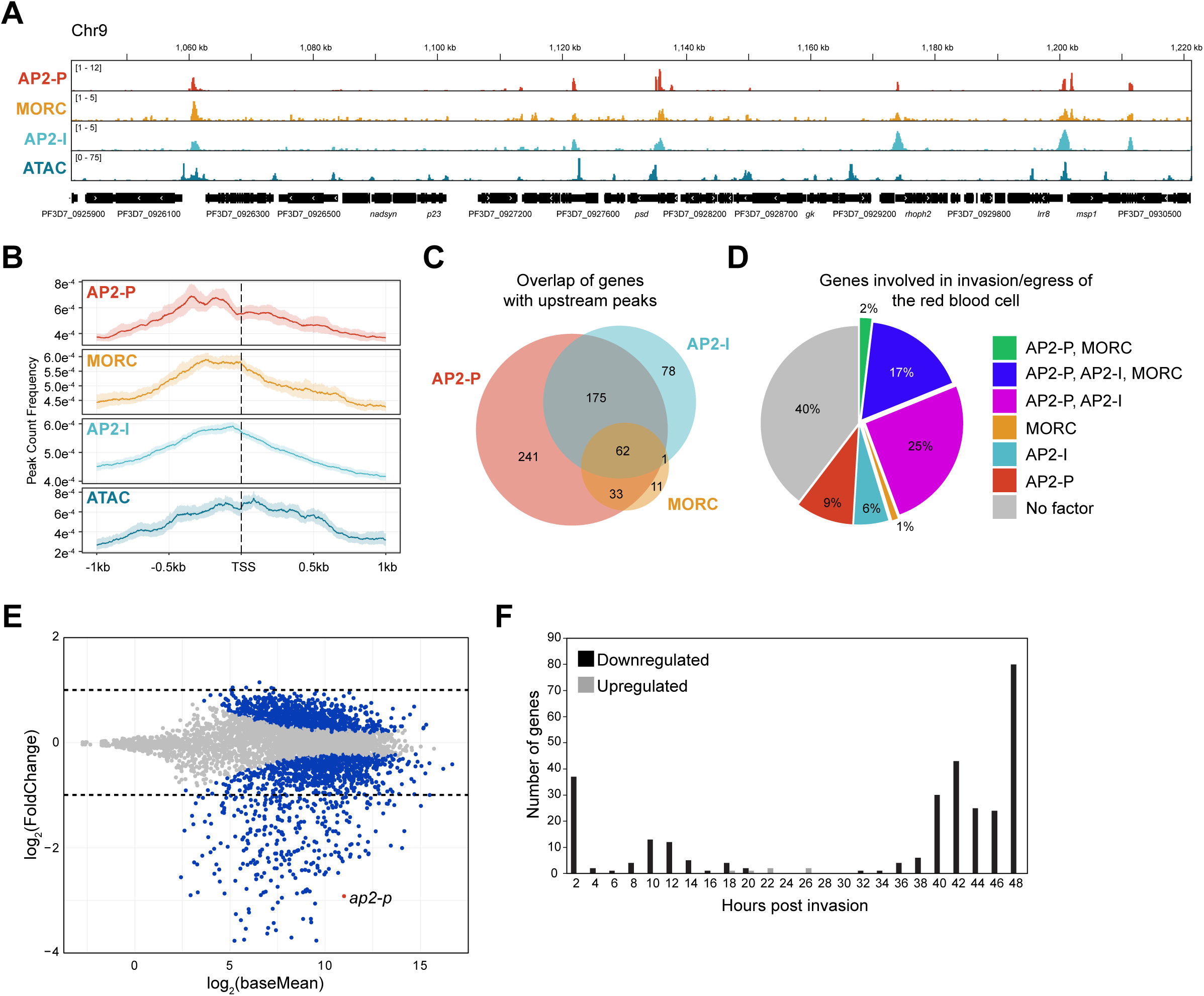
AP2-P, AP2-I, and MORC activate stage-specific genes. **(A)** ChIP-seq data showing enrichment (*y*-axis = ChIP/Input) of AP2-P, MORC, and AP2-I^62^ in clonal late-stage parasites at a central region of chromosome 9. ATAC-seq data^58^ from a closely corresponding stage showing chromatin accessibility [*y*-axis = ATAC-seq (RPM)/gDNA (RPM)]. The *x*-axis is DNA sequence, with genes represented by black boxes with white arrowheads to indicate transcription direction. **(B)** Metagene plots showing average AP2-P and MORC enrichment in clonal AP2-P-3HA-*glmS* and MORC-3HA parasites, respectively, in late-stage parasites from 1 kb upstream to 1 kb downstream of the TSS of all genes with an upstream peak of AP2-P, excluding *var* genes. AP2-I ChIP-seq data^62^ and ATAC-seq data^58^ from stage-matched parasites is included. One replicate was used for the AP2-P and MORC ChIP datasets. **(C)** Venn diagram showing overlap of genes bound by AP2-P, MORC, and AP2-I in their upstream regions (**Table 1**). **(D)** Pie charts showing the proportion of genes that are important to the late stages of the parasite asexual replication cycle^64,65^ (**Table 4**) that are bound by AP2-P, MORC, and/or AP2-I in their upstream regions (**Table 1**). **(E)** Differential expression analysis of AP2-P KD. MA plot of log_2_(knockdown/control, M) plotted over the mean abundance of each gene (A) in late-stage parasites. Transcripts that were significantly higher (above *x*-axis) or lower (below *x*-axis) in abundance after AP2-P KD are highlighted in blue (*q* ≤ 0.05). *ap2-p* is highlighted in red. Dotted lines indicate a fold change ≥ 2. Three technical replicates were used for untreated and glucosamine-treated parasites. P-values were calculated with a Wald test for significance of coefficients in a negative binomial generalized linear model as implemented in DESeq2^108^. *q* = Bonferroni corrected P-value. **(F)** Frequency plot showing the time in the red blood cell cycle (hours post invasion of the red blood cell) of peak transcript level (comparison to transcriptomics time course in^63^) for genes that are significantly downregulated (black) or upregulated (grey) more than two-fold following AP2-P knockdown in late-stage parasites.

Gene Ontology (GO) analysis of genes with an upstream AP2-P peak showed that the most significantly enriched terms were “cell-cell adhesion” (*q* = 3.95 x 10^-11^) and “entry into host” (*q* = 1.18 x 10^-9^), categories which include *var* genes as well as genes involved in red blood cell invasion (**Table 3**). While *var* genes are silent in late-stage parasites, genes involved in red blood cell egress and invasion are specifically and highly transcribed at this stage^63^. Indeed, approximately 60% of a cohort of genes whose role in red blood cell egress and invasion have been well characterized^64,65^ (**Table 4**) are bound by combinations of AP2-P, AP2-I, and MORC in their upstream regions (**Fig. 4D**).

To determine if AP2-P plays a role in transcriptional regulation of late-stage-specific genes, we revisited the AP2-P KD differential expression analysis. AP2-P KD resulted in the significant upregulation of only six genes, but downregulation of 302 genes more than two-fold, suggesting that AP2-P plays a transcriptional activating role at this stage (**Fig. 4E**; **Table 2**). Downregulated genes are most enriched for GO terms such as “movement in host environment” (*q* = 1.04 x 10^-22^) and “entry into host” (*q* = 9.63 x 10^-22^) (**Table 5**), and most of them normally reach peak transcription levels in late-stage parasites^63^ (**Fig. 4F**). Moreover, downregulated genes are enriched in AP2-P, MORC, and AP2-I in their upstream regions (**Supp.** Fig. 4B; **Table 1**). Taken together, these data suggest that AP2-P, AP2-I, and MORC form an activating complex within euchromatic regions that is essential for the transcription of genes that are specific and important during late stages of the red blood cell cycle.

### AP2-P activating complex defines euchromatic boundaries and long-range interactions amongst functionally related genes

Given that AP2-P and MORC define the boundaries of subtelomeric heterochromatic self-associating domains, we investigated whether they are also involved in the formation of boundaries of euchromatic genome structures. In euchromatic regions of the parasite genome, we observed significant breaks in local contacts, or boundaries, that often overlap with genes showing high levels of transcription, such as *dblmsp* (PF3D7_1035700) and *gap45* (PF3D7_1222700) (**Fig. 5A,B**; **Table 6**). Indeed, genes near these euchromatic boundaries have higher levels of transcription than those outside of boundaries (**Fig. 5C**), and euchromatic boundary strength shows linear correlation to transcription levels of nearby genes (**Supp.** Fig. 5A). Importantly, euchromatic boundaries show significant overlap with genes associated with AP2-P, AP2-I, and/or MORC peaks (**Fig. 5A,D**; **Supp.** Fig. 5B). In fact, we observed a positive correlation between boundary insulation score and enrichment of AP2-P, AP2-I, and MORC (**Fig. 5D**). This pattern contrasts with that seen at non-genic subtelomeric boundaries, where insulation score correlates with enrichment of AP2-P, TRZ, and MORC (**Fig. 5D**). These data suggest that different protein complex compositions may define different types of boundaries.

**Figure 5:**
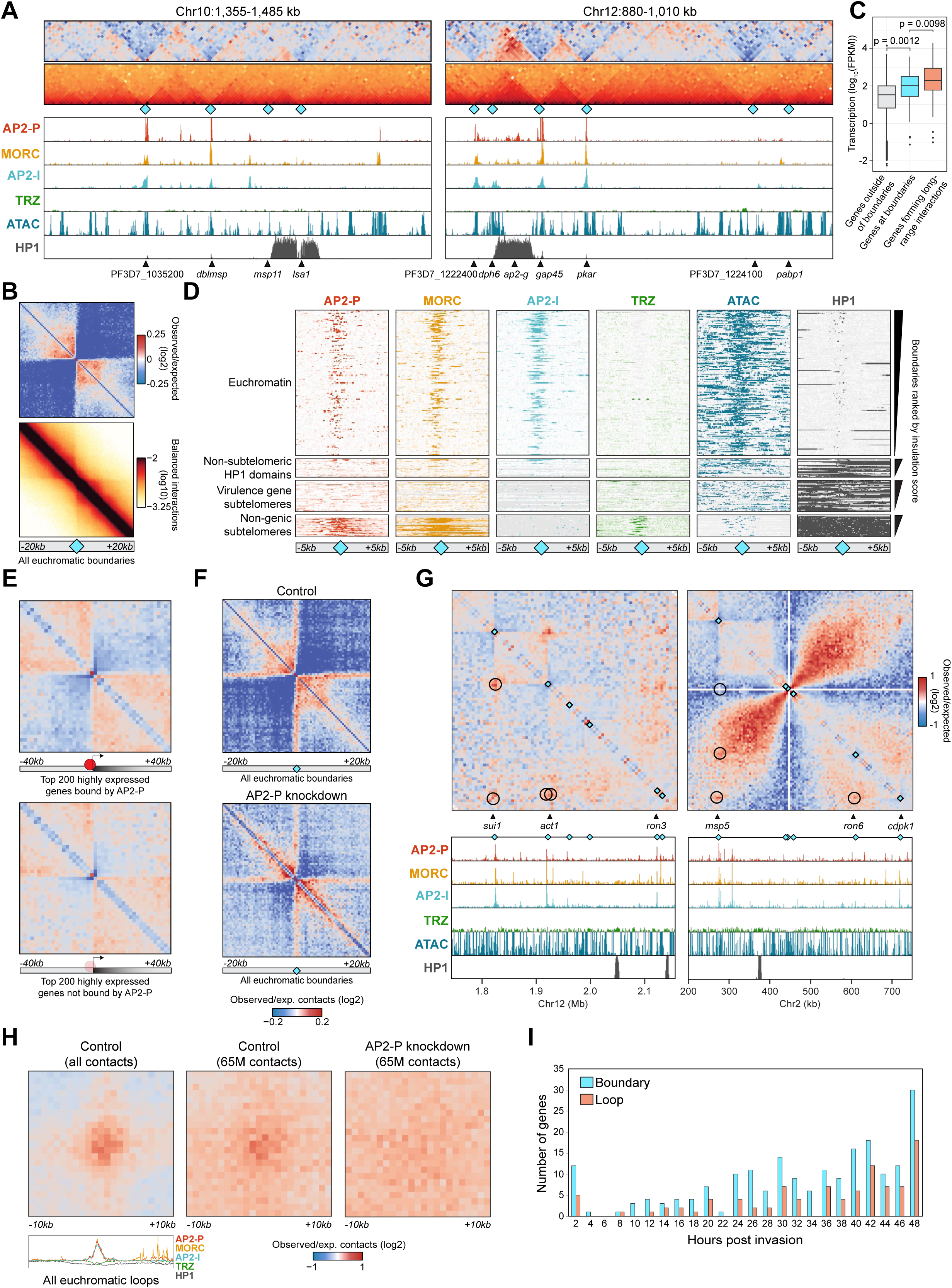
AP2-P activating complex defines euchromatic boundaries and long-range interactions amongst functionally related genes. **(A)** Micro-C contact maps of central regions of chromosome 10 (left) and 12 (right) in late-stage parasites (130kb-wide loci, 1kb resolution). Micro-C boundaries are indicated with cyan diamonds. Color scales indicate log_2_-scaled observed/expected interaction frequency ratios (top) and normalized interaction frequency (bottom). Corresponding AP2-P, MORC, AP2-I^62^, TRZ^60^, and HP1^110^ ChIP/Input ratio and ATAC-seq^58^ data from late-stage parasites are shown below. Selected genes are indicated at the bottom. **(B)** On-diagonal Micro-C contact map, aggregated over the 211 euchromatic boundaries (indicated with cyan diamond, **Table 6**) identified by chromosight^94^ (±20kb, 500bp resolution). Color scales indicate log_2_-scaled observed/expected interaction frequency ratios (top) and normalized interaction frequency (bottom). **(C)** Transcript levels (log10FPKM) of genes outside of Micro-C boundaries (grey), associated with a euchromatic Micro-C boundary (cyan, **Table 6**), or associated with a euchromatic long-range looping event (orange, **Table 7**) (see Methods). Statistical difference in levels of expression between each set of genes was assessed using t-tests. **(D)** Heatmap showing AP2-P, MORC, AP2-I^62^, TRZ^60^, and HP1^110^ enrichment or ATAC-seq signal^58^ around 317 boundaries identified by chromosight^94^ (± 5kb, shown below with cyan diamonds, **Table 6**). Rows represent Micro-C boundaries ranked by insulation score in euchromatin, non-subtelomeric HP1-enriched domains, virulence gene-containing subtelomeric regions, or non-genic subtelomeric regions. The colors indicate the normalized ChIP-seq/ATAC-seq enrichment level. **(E)** On-diagonal Micro-C contact maps, aggregated over the TSS of the 200 genes most highly expressed in late-stage parasites (±40kb, 2kb resolution) and associated with AP2-P (≤ 1kb from an AP2-P peak) or not. Color scales indicate log_2_-scaled observed/expected interaction frequency ratios. **(F)** On-diagonal Micro-C contact maps, aggregated over all 211 euchromatin boundaries (cyan diamonds, ±20kb, 500bp resolution, **Table 6**). Color scales indicate log_2_-scaled observed/expected interaction frequency ratios in subsampled (65 million contacts) control map (top) or AP2-P KD map (bottom). **(G)** Micro-C contact map of central regions of chromosome 12 (left) and 2 (right) in late-stage parasites. Micro-C boundaries are indicated with cyan diamonds. Selected genes are indicated below. AP2-P, MORC, AP2-I^62^, TRZ^60^, and HP1^110^ ChIP/Input ratio signals and ATAC-seq^58^ data from late-stage parasites are shown at the bottom. Long-range loops identified with chromosight^94^ are indicated with black circles on the contact map. **(H)** Off-diagonal Micro-C contact maps, aggregated over the 85 euchromatin loops (±10kb, 1kb resolution) found in late-stage parasites. Color scales indicate log_2_-scaled observed/expected interaction frequency ratios in the control map (left), subsampled control map (center), or AP2-P KD map (right). 1D aggregated AP2-P, MORC, AP2-I^62^, TRZ^60^, and HP1^110^ ChIP/input signals from late-stage parasites are shown below. (I) Frequency plot showing the time in the red blood cell cycle (hours post invasion of the red blood cell) of peak transcript level (comparison to transcriptomics time course in^63^) for genes that are associated with a euchromatic boundary (cyan) or looping event (orange) in late-stage parasites.

We further found that AP2-P-bound active genes have a higher boundary insulation score than genes with similar levels of transcription that are not bound by AP2-P (**Fig. 5E**). Moreover, AP2-P KD leads to weakening of euchromatic boundaries (**Fig. 5F**). These data suggest that strong transcriptional activity could play a role in boundary formation, as seen in other microorganisms^17,18^, and that AP2-P directly contributes to the structural maintenance of euchromatic boundaries.

Interestingly, we observed several long-range intra- and interchromosomal contacts involving multiple active genes across the genome (**Table 7**). Such intrachromosomal long-range interactions can be seen between *sui1* (PF3D7_1243600) and two other genes on chromosome 12: *act1* (PF3D7_1246200, separated by ∼94kb) and *ron3* (PF3D7_1252100, separated by ∼300kb) (**Fig. 5G**, left). Another example can be found on chromosome 2, where *msp5* (PF3D7_0206900) interacts with *ron6* (PF3D7_0214900, separated by ∼334kb) and *cdpk1* (PF3D7_0217500, separated by ∼445kb) (**Fig. 5G**, right). These contacts even span the centromere on chromosome 2. One final example is on chromosome 10 where *chd1* (PF3D7_1023900) interacts with *nprx* (PF3D7_1027300, separated by ∼134kb), *msp11* (Pf3D7_1036000, separated by ∼ 420kb), and PF3D7_1035200 (separated by ∼384kb), which in turn interacts with *etramp10.2* (PF3D7_1033200) (**Supp.** Fig. 5C). Several of the above-mentioned loci on different chromosomes also form trans contacts with each other, such as *msp5* on chromosome 2, which interacts with multiple loci on chromosome 12 –*sui1* and *act1* (**Supp.** Fig. 5D).

Importantly, the loci that form these long-range contacts are enriched in AP2-P, AP2-I, and MORC (**Fig. 5G,H left**; **Supp.** Fig. 5C). Furthermore, AP2-P KD reduced contact frequency between these loci (**Fig. 5H**, middle and right). Finally, genes that form these long-range interactions are transcribed at levels even higher than those simply associated with euchromatic boundaries (**Fig. 5C**) and have characteristics typical of late stage-specific genes: they are most significantly enriched in the GO term “entry into host” (*q* = 4.32 x 10^-8^, Table 8) and tend to reach peak transcription levels in late-stage parasites (**Fig. 5I**). Taken together, these data suggest that AP2-P (and perhaps MORC and AP2-I) facilitate long-range interactions between highly transcribed, stage-specific genes.

## Discussion

Here we adapted the Micro-C technique^18^ to the human malaria parasite *P. falciparum* to map genome-wide contacts at near-nucleosome resolution. With this increased resolution, we were able to define two different types of heterochromatin: HP1-enriched heterochromatin encompassing *ap2-g* or virulence genes (such as the *var* family) is well insulated from neighboring euchromatin and forms significant intra- and interchromosomal interactions, whereas other genes enriched in HP1 do not show these same characteristics, suggesting that other as-yet unidentified protein factors are involved in insulation of and contacts amongst *ap2-g* and virulence genes (**Fig. 6**, top). In addition, we identified two new chromatin structures in blood stage parasites. First, we discovered heterochromatic subtelomeric fold structures that are distinct from telomere loops, which protect the ends of chromosomes (**Fig. 6**, bottom left).

**Figure 6:**
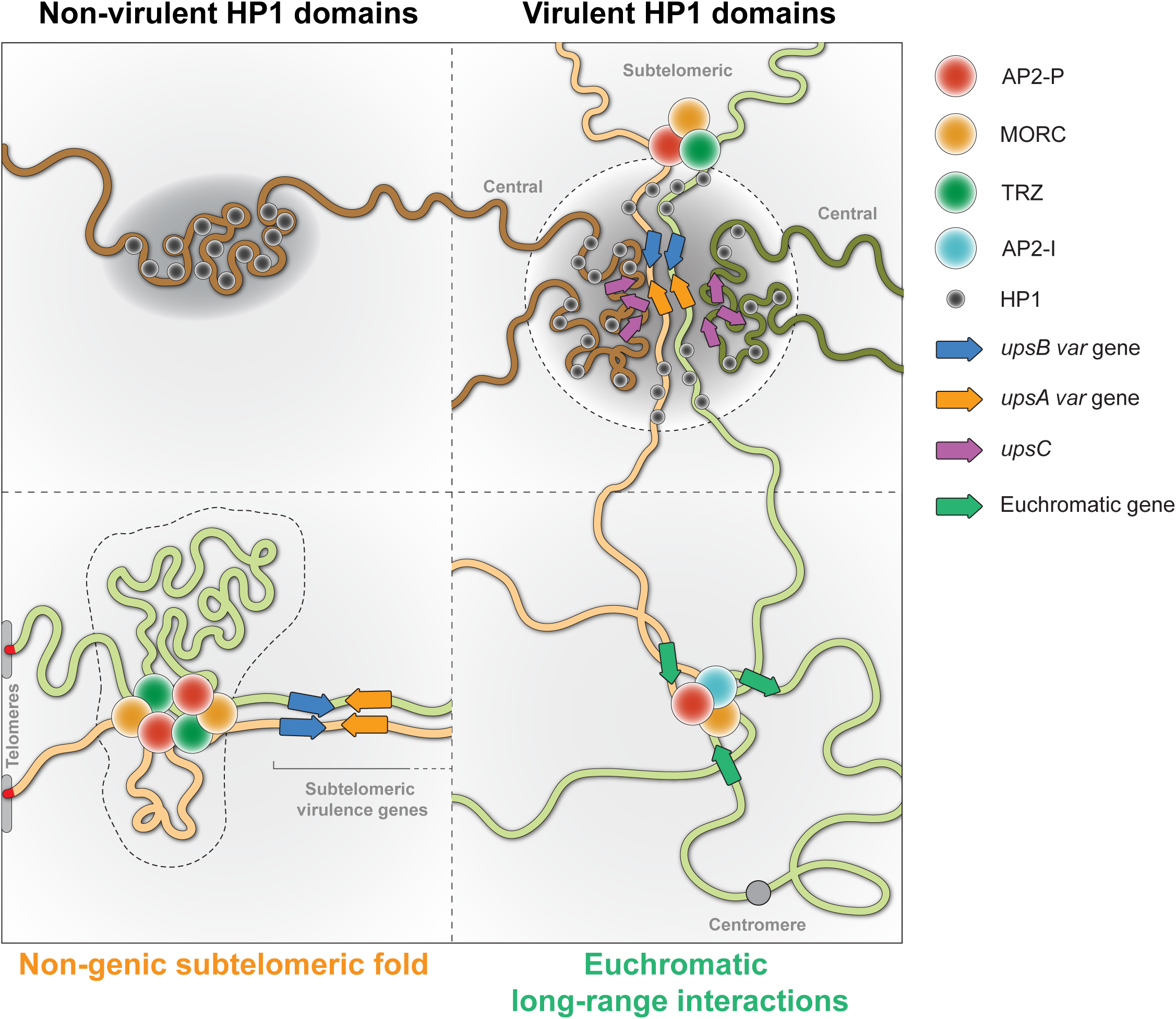
Schematic of heterochromatic and euchromatic structures observed with Micro-C.

These subtelomeric folds could play a role in coordinating *var* gene alignment. Second, we observed long-range intra- and interchromosomal interactions amongst stage-specific genes, which could facilitate their temporal co-activation (**Fig. 6**, bottom right). Integration of Micro-C data, ChIP- and RNA-seq data, and functional gene characterization identified the proteins responsible for forming these two different higher order structures: AP2-P and MORC.

### AP2-P and MORC form subtelomeric structures to facilitate *var* gene alignment

AP2-P and MORC are most significantly enriched upstream of subtelomeric *var* genes at a clear Micro-C boundary between the HP1-enriched non-genic and genic domains, which is also enriched in TRZ and a second ApiAP2 DNA binding protein, SIP2. This site is 1) the anchor point of a fold structure formed by the non-genic subtelomeric region and 2) the main point of contact between subtelomeric regions of the same and different chromosomes. Both types of structures are disrupted upon AP2-P KD, suggesting a direct role in structure formation. Our data suggest that while HP1 is a mediator of subtelomeric heterochromatinization, AP2-P, MORC, and perhaps other proteins such as TRZ and SIP2 play a crucial role in forming focal contacts within and between chromosomes at the heterochromatic boundary between the non-genic and genic subtelomeric regions. This conclusion is supported by two recent studies that used Hi-C to show a decrease in virulence gene clustering upon knockout of AP2-P^66^ or KD of MORC^67^. Since strong contacts amongst telomeres facilitate chromosome end clustering, the additional subtelomeric points of contact mediated by AP2-P/MORC could play a role in *var* gene biology.

Because the level to which we knocked AP2-P down did not result in death or cell cycle arrest, we were able to gain clear insight into its chromatin-related function in late-stage parasites. Importantly, we did not observe a significant change in *var* gene transcription upon AP2-P depletion, as observed elsewhere^66^, which is in line with the fact that AP2-P and MORC do not bind to the *var* gene promoter, but considerably farther upstream (**Fig. 2C**). Thus, we hypothesize that AP2-P/MORC-mediated DNA-DNA interactions rather play a role in *var* gene genetics. The subtelomeric *upsB var* genes in particular are known to undergo ectopic recombination during mitosis^42,43^ that leads to antigenic diversity, which is crucial to maintaining chronic infection in the human host. Unlike V(D)J recombination amongst immunoglobulin genes^68^, *var* genes are not targeted for recombination by a specific homologous sequence motif; in fact, *var* gene recombination does not rely on homology^43^. To recombine in a manner that preserves sequence integrity, *var* genes would presumably need to be brought into close proximity to one another via some other means.

While telomeres cluster at the nuclear periphery, subtelomeric *var* genes are located at vastly different distances from the chromosome end depending on the chromosome (mean 26kb ± 11.9kb). Regardless of the length of the non-genic subtelomeric region, AP2-P and MORC form a subtelomeric fold specifically anchored ∼8kb downstream of the telomere (mean 8.3kb ± 1.7kb) and ∼2kb upstream of the first *var* gene on the chromosome end (mean 2.1kb ± 0.22kb) (**Fig. 2B,C**). Consistent with its ability to topologically entrap DNA and multimerize in other eukaryotes^69^, *Pf*MORC could bind different genomic loci, dimerize, and cinch up the non-coding subtelomeric sequence in a way that brings the first *var* gene on the chromosome end close to the telomere. Then, AP2-P, MORC, and SIP2 or TRZ could facilitate intra-and interchromosomal subtelomeric interactions to bring *var* genes in line with each other. We propose a model in which AP2-P and MORC facilitate subtelomeric fold structures to cluster and align *var* genes in space so that they can recombine properly (**Fig. 6, bottom left**).

### AP2-P and MORC activate genes in a spatio-temporal manner

In addition to their role in subtelomeric heterochromatin structure, we also show that AP2-P and MORC play a role in euchromatin structure and gene activation. These factors show synergism with AP2-I at the promoters of active, late-stage-specific genes that form intra- and interchromosomal long-range contacts (**Fig. 6, bottom right**). Long-range interactions between genes have been observed in multiple different eukaryotes and can be associated with co-repression or co-expression. In humans, long-range looping interactions, often mediated by cohesin and CTCF, between cis regulatory elements (such as enhancers and promoters) can activate genes in specific cellular or developmental contexts^70–72^. In contrast, the long-range euchromatic contacts we observe do not seem to involve an enhancer element or cohesin^49^, and *P. falciparum* does not have CTCF^44^. AP2-P/MORC-mediated euchromatic long-range contacts more closely resemble the interactions seen between co-expressed and/or functionally related genes in *Drosophila* chromosome conformation capture data^9,73,74^. A recent study provided functional evidence that looping between distant genes helps to coordinate and fine-tune their expression during *Drosophila* embryonic development^74^. We propose a similar mechanism exists in *P. falciparum* between stage-specific genes.

It has been suggested that transcription does not happen from individual promoters in a random and dispersed manner in the nucleus, but rather in foci where the transcriptional machinery is concentrated^23,24^. Some have hypothesized that in mammals, these condensates represent phase separated compartments where multiple gene promoters can associate with enhancers that lead to “bursts” of transcription^75,76^. In other eukaryotes, highly active genes have been shown to associate with compartments in the nucleus such as nuclear speckles, which have high concentrations of factors involved in transcription and transcript processing^77^. Such an “activation center” has been proposed to exist for the activation of a single member of the *var* multigene family in *P. falciparum*^78^ and the variant surface glycoprotein multigene family in *Trypanosoma brucei*^79,80^. We have shown that in earlier stages of the red blood cell cycle, a component of the cohesin complex binds to and helps repress the same cohort of genes bound and activated by AP2-P, MORC, and AP2-I in late-stage parasites^49^. Thus, AP2-P/MORC binding to these genes in late-stage parasites may dislodge the cohesin complex and re-locate them to an “activation center”.

Although the exact biophysical nature of long-range DNA-DNA contacts in *P. falciparum* remains to be determined, our data suggest that they are dependent on the site-specific interaction of AP2-P and MORC. Spatial relocation of genes to compartments where the transcriptional machinery is concentrated may facilitate rapid activation, or transcriptional bursts, in a stage-specific manner. Association of functionally related genes has also been observed in sporozoites, the stage that is injected by the mosquito into the human host^37^. Thus, spatio-temporal regulation of stage-specific genes may be crucial to driving the entire parasite life cycle. Further investigation is needed of the properties, dynamics, and significance of the coalescence of co-regulated genes.

### AP2-P and MORC serve as versatile genome organizing factors

By characterizing long-range heterochromatic and euchromatic DNA-DNA interactions, our study has highlighted the versatility of AP2-P and MORC. Among the Apicomplexan parasites studied, MORC has been observed via mass spectrometry to interact with many different ApiAP2 factors^50,55,66,67,81–83^, yet the nature and function of these interactions was unclear in *Plasmodium*. We now provide genome-wide evidence for the functional cooperation between MORC and three ApiAP2 factors – AP2-P, SIP2, and AP2-I – at the chromatin level. As MORC in other eukaryotes does not seem to have DNA binding specificity, it is likely that *Pf*MORC is an effector protein that is targeted to specific loci by multiple ApiAP2 factors. Indeed, cooperative binding of two different ApiAP2 factors and MORC plays an important role in the repression of stage-specific genes in a related apicomplexan parasite, *Toxoplasma gondii*^55,83^.

Combinatorial binding amongst ApiAP2 factors and their effector proteins, such as MORC, could therefore provide a higher capacity for complex transcriptional regulation than a “one transcription factor:one gene” model, especially in this organism that has relatively few sequence-specific DNA binding factors^46^. In this regard, the function of a single ApiAP2 factor – AP2-G – as a master inducer of sexual commitment (gametocytogenesis) might be an exception rather than the rule^84,85^.

While MORC is associated with heterochromatin and transcriptional repression in other eukaryotes^51^, including the apicomplexan parasite *T. gondii*^55^, we and others^50,66,67,82^ find AP2-P and MORC enrichment at silent *and* active genes. It is likely that this versatile binding results from its association with different DNA-binding factors such as SIP2 and AP2-I, which are more limited in their binding to heterochromatic and euchromatic regions, respectively. Therefore, it is possible that other ApiAP2 factors that show peak abundance at other stages of the parasite life cycle can equally recruit MORC to their targets in a stage-specific manner.

Whether in eu- or heterochromatin, AP2-P and MORC are enriched at anchor points for long-range DNA-DNA interactions. We therefore propose a unifying model where AP2-P and MORC facilitate higher order chromatin structure, which influences different context-specific chromatin-associated processes such as recombination and transcription. In this way, MORC and AP2-P may replace organizational proteins such as CTCF, which *Plasmodium* lacks. Our study of genome organization in a eukaryotic parasite with a massive global health impact has shed light on divergent and potentially targetable molecular mechanisms used to achieve similar complex phenomena observed in model eukaryotes: spatial co-regulation of functionally related genes.

## Acknowledgements

This work was supported by the Agence Nationale de la Recherche (grants ANR-21-CE15-0002-02 ApiMORCing and ANR-21-CE15-0010-01 PlasmoVarOrg to J.M.B.); the European Research Council (grant PlasmoEpiRNA 947819 to S.B. and ERC grant agreement 771813 to R.K.); Emerging Infectious Diseases junior seed grant from the Pasteur Institute to J.C. and C.R. P.S. was supported by the Institut Pasteur Roux-Cantarini postdoctoral fellowship. J.S. was supported by a postdoctoral fellowship from l’Association pour la Recherche sur le Cancer (ARC). The authors would like to acknowledge the use of the Biomics and Flow Cytometry platforms at the Institut Pasteur and the invaluable support of PlasmoDB and the ParaFrap network.

## Author Contributions

Conceptualization, J.M.B., S.B.; Methodology (Micro-C adaptation to *P. falciparum*), J.C., J.M.B., S.B.; Formal analysis, P.S., J.S., S.B.; Investigation, J.M.B., P.S., J.C., M.M.C., C.R., P.C.; Visualization, J.M.B., P.S., J.S.; Writing – Original Draft, J.M.B.; Writing – Review & Editing, all authors; Funding Acquisition, J.M.B., S.B., R.K.; Supervision, J.M.B., S.B., R.K., A.S.

## Declaration **of Interests**

The authors declare no competing interests.

## Methods

### Parasite culture

Blood stage 3D7 or NF54 *P. falciparum* parasites were cultured as previously described^36^. Briefly, parasites were cultured in human RBCs supplemented with 10% v/v Albumax I (Thermo Fisher 11020), hypoxanthine (0.1 mM final concentration, C.C.Pro Z-41-M) and 10 mg gentamicin (Sigma G1397) at 4% hematocrit and under 5% O2, 5% CO2 at 37 °C. Parasites were synchronized by sorbitol (5%, Sigma S6021) lysis during ring stage followed by a plasmagel (Plasmion, Fresenius Kabi) enrichment for late blood stages 24 hours later. Another sorbitol treatment 6 h afterwards places the 0 hpi (hours post invasion of the red blood cell) time point 3 h after the plasmagel enrichment. Thus, the window of synchronicity for cultures is ± 3 h. In this manuscript, “early-stage”, “middle-stage”, and “late-stage” refers to 12, 24, and 36 hpi ± 3 h, respectively. Parasite development was monitored by Giemsa staining. Parasites were harvested at 1–5% parasitemia.

### Micro-C

The Micro-C protocol was performed as previously described^18^. NF54 WT parasites were synchronized, and eight replicates (∼8 x 10^8^ schizonts each) were lysed with saponin (0.075% in DPBS) and washed with DPBS at 37°C. Parasites were resuspended in DPBS at 25°C and cross-linked for 10 min by adding methanol-free formaldehyde (ThermoFisher 28908) to 1% final concentration with gentle agitation. The reaction was quenched by adding 1 M Tris-HCl pH 7.5 to a final concentration of 0.75 M and incubating at 25°C for 5 min with gentle agitation.

Parasites were centrifuged for 5 min at 3,250g, washed with DPBS at 25°C, resuspended in the second cross-linking solution - 3 mM DSG (ThermoFisher 20593) in DPBS – and incubated for 45 min at 25°C with agitation. The reaction was quenched with 1 M Tris-HCl pH 7.5 to a final concentration of 0.75 M and incubated at 25°C for 5 min with agitation. The double cross-linked parasites were washed with DPBS at 25°C, and the pellets were snap-frozen and stored at −80 °C until further use.

One replicate was split into four and used to titrate the MNase (Worthington Biochem LS004798). Thus, 4 x ∼2 x 10^8^ parasites were resuspended in 1 ml of MB#1 [50 mM NaCl, 10 mM Tris-HCl pH 7.5, 5 mM MgCl_2_, 1 mM CaCl_2_, 0.2% NP-40, 1x protease inhibitor (PI, Roche 4693159001)] and incubated on ice for 20 min with regular flicking. The cells were centrifuged (10,000 *g*, 5 min, 4 °C), washed with MB#1 and resuspended in 190 μl of MB#1. 10 μl of MNase at the chosen concentration (corresponding to 10U, 20U, 50U and 100U) was added, and the mixture was incubated for exactly 10 min at 37°C with continuous shaking at 850 rpm. To stop the reaction, 500 mM EGTA (ThermoFisher J60767) was added to a final concentration of 4 mM and was followed by a 10 min incubation at 65°C. The digested nuclei were then washed twice (5 min centrifugation at 16,000 *g*, 4°C) with MB#2 (50 mM NaCl, 10 mM Tris-HCl pH 7.5, 10 mM MgCl_2_). The nuclei were resuspended in a digestion mix [10 mM Tris-HCl pH 7.5, 1 mM EDTA, 1% SDS, 1 mg/ml Proteinase K (ThermoFisher EO0491), 0.125 mg/ml RNase A (ThermoFisher EO0491)] and decrosslinked for 10 h at 65°C. The samples were centrifuged (16,000 *g*, 10 min, 4°C) and DNA contained in the supernatant was extracted using the DNA Clean & Concentrator kit (Zymo D4013). The fragment sizes were assessed by Agilent BioAnalyzer to choose the appropriate MNase concentration yielding a 90/10 monomer/dimer ratio.

The following steps were performed with four replicates. The cell pellet (∼8 x 10^8^ parasites) was lysed and digested with the chosen concentration of MNase, as described above. After the MB#2 washes, each pellet was centrifuged (16,000 *g*, 5 min, 4°C), resuspended in 45 μl of end-chewing mix [1X NEB Buffer 2.1, 2 mM ATP (ThermoFisher R1441), 5 mM DTT (ThermoFisher 707265ML), 0.5 U/μl T4 PNK (NEB M0201S)] and incubated at 37°C for 15 min on an agitator with interval mixing (800 rpm 10 s, resting 30 s). Five μl of 5 U/μl Klenow fragment (NEB M0210L) was added to achieve a final concentration of 0.5 U/μl and incubated at 37°C for 15 min with the same agitation program as the previous step. 25 μl of end-labeling mix [0.2 mM Biotin-dATP (Jena Bioscience NU-835-BIO14-S), 0.2 mM Biotin-dCTP (Jena Bioscience NU-809-BIOX-S), 0.2 mM dTTP (ThermoFisher 18255018), 0.2 mM dGTP (ThermoFisher 18254011), 0.1 mg/ml BSA (Invitrogen AM2616)]] was added to the mixture and incubated at 25°C for 45 min with agitation (800 rpm 10 s, resting 30 s). 500 mM EDTA (Invitrogen AM9260G) was added to a final concentration of 30 mM and incubated for 30 min at 65°C. The pellet was centrifuged (10,000 *g*, 5 min, 4°C) and washed with cold MB#3 (10 mM Tris-HCl pH 7.5, 10 mM MgCl_2_). The biotinylated DNA fragments were then covalently linked by proximity ligation by resuspending the pellet in 250 μl of ligation mix [1X T4 DNA ligase buffer (NEB B0202S), 0.1 mg/ml BSA (Invitrogen AM2616), 33 U/μl T4 DNA ligase (NEB M0202L)] and incubating at 25°C for 2.5 h with slow rotation. The ligated DNA fragments were then centrifuged (16,000 *g*, 5 min, 4°C), and the biotin-dNTPs were removed from unligated ends by resuspending the pellet in 100 μl of 1X NEB buffer #1 (NEB B7001S) and 5 U/μl exonuclease III (NEB M0206L) and incubating for 15 min at 37°C with agitation (800 rpm 10 s, resting 30 s).

The DNA was purified. 12.5 μl of 10% SDS (ThermoFisher 15553027) was added, and the DNA was decrosslinked by incubating for 10 h at 65°C. RNase A (ThermoFisher EN0531) was added to a final concentration of 0.2 mg/ml and incubated for 2 h at 37°C. Proteinase K (ThermoFisher EO0491) was added to a final concentration of 0.2 mg/ml and incubated for 2 h at 55°C. After centrifugation (16,000g, 10 min, 4 °C), the DNA contained in the supernatant was recovered using the DNA Clean & Concentrator (Zymo D4013, using a 5:1 ratio of DNA Binding Buffer to Sample). The ligated DNA was mixed with loading dye (ThermoFisher R1161) and loaded onto a 2% TAE agarose gel to separate dimers from monomers. The gel was run at 100V until a satisfying separation was obtained. A band from 250-400bp was cut and DNA was extracted in 150 μl of ddH2O using the Gel DNA Recovery Kit (Zymo D4007).

The biotinylated dimers were isolated with Dynabeads MyOne Streptavidin C1 (ThermoFisher 65001). The beads were washed with 500 μl TBW (1 M NaCl, 5 mM Tris-HCl pH 7.5, 0.5 mM EDTA, 0.10% Tween20) on a magnet and resuspended in 150 μl 2X BW (2 M NaCl, 10 mM Tris-HCl pH 7.5, 1 mM EDTA). The ligated fragments were added and rotated for 20 min at 25°C. The beads were then washed twice with 500 μl TBW at 55°C on an agitator (2 min, 800 rpm 10 s, resting 30 s) and once with EB (10 mM Tris-HCl pH 7.5) at 25°C. The library prep was performed with the samples on beads using the NEBNext® Ultra II Directional DNA Library Prep Kit (NEB E7645), with several modifications. First, the end repair and adapter ligations steps were performed with interval mixing (800 rpm 10 s, resting 30 s). After the adapter ligation, the beads were washed twice with TWB at 55°C and once with EB (as described above) and resuspended in 20 μl EB for the PCR reaction. No size-selection step was performed. Finally, after the PCR, the beads were placed again on a magnet, and the supernatant was purified with AMPure beads as recommended.

This protocol was repeated for the AP2-P KD in the AP2-P-3HA strain (clone H5 in 3D7 background). AP2-P-3HA-*glmS* (clone H5) parasites were synchronized, and the culture was split into two at 12 hpi. Glucosamine (Sigma G1514, final concentration 2.5 mM) was added to one culture for two rounds of parasite replication (approximately 96 h). Parasites were then re-synchronized and four technical replicates (with and without glucosamine) were harvested at 36 hpi.

### Generation of strains

The AP2-P-3HA-*glms* strain was generated using a two-plasmid system (pUF1 and pL7) based on the CRISPR/Cas9 system previously described^86^. A 3D7 wild-type bulk ring stage culture was transfected with 25 μg pUF1-Cas9 and 25 μg of pL7-PF3D7_1107800-3HA-*glmS* containing a single guide RNA (sgRNA)-encoding sequence targeting the 3’ UTR of PF3D7_1107800 (Table 9). The pL7-PF3D7_1107800-3HA-*glmS* plasmid also contained a homology repair construct synthesized by GenScript Biotech (Table 9). This homology repair construct comprises a 3 x Hemaglutinin (3HA)-encoding sequence followed by a *glmS* ribozyme-encoding sequence^61^, which are flanked by ∼300 bp homology repair regions upstream and downstream of the Cas9 cut site. Two shield mutations were made in the upstream homology repair region to prevent further Cas9 cleavage of the modified locus. The sgRNA sequence was designed using Protospacer^87^. The sgRNA sequence uniquely targeted a single sequence in the genome. After transfection, drug selection was applied for five days at 2.67 nM WR99210 (Jacobus Pharmaceuticals) and 1.5 μM DSM1 (MR4/BEI Resources). Parasites reappeared approximately three weeks after transfection, and 5-fluorocytosine was used to negatively select the pL7 plasmid.

The MORC-3HA and AP2-P-3HA-*glmS*:MORC-GFP strains were generated using the method of selection-linked integration (SLI) previously described^88^, with a slight modification wherein a GSG-encoding sequence (5’ – GGTAGTGGT – 3’) was added directly upstream of the T2A skip peptide-encoding sequence to enhance cleavage of the tagged protein from the downstream drug selection protein. The homology region corresponding to the 771 bp at the 3’ end of *morc* (PF3D7_1468100) was amplified using JB_155F/R (Table 9). The PCR fragment was then fused to a sequence encoding a 3HA or GFP epitope tag followed by a skip peptide followed by the neomycin resistance marker. For the MORC-3HA strain, a 3D7 wild-type bulk ring stage culture was transfected with 50 μg of pSLI plasmid (containing a yeast dihydroorotate dehydrogenase selection marker) and selected first with 1.5 μM DSM1 (MR4/BEI Resources). Parasites reappeared approximately three weeks after transfection and positive selection for integration was performed via the addition of 400µg/mL G418 (Sigma G8168). For the AP2-P-3HA-*glmS*:MORC-GFP strain, a clonal (clone H5) culture of AP2-P-3HA-*glmS* was transfected at ring stage with 50 μg of pSLI plasmid (containing a blasticidin selection marker) and selected first with 5 μg/mL blasticidin (Invivogen ant-bl-05). Parasites reappeared approximately five weeks after transfection and positive selection for integration was performed via the addition of 400µg/mL G418 (Sigma G8168).

All cloning was performed using KAPA HiFi DNA Polymerase (Roche 07958846001), In-Fusion HD Cloning Kit (Clontech 639649), and XL10-Gold Ultracompetent *E. coli* (Agilent Technologies 200315). All the parasite lines were cloned by limiting dilution, and integration at the targeted genomic locus was confirmed by PCR (**Supp.** Fig. 2D,E; **Supp.** Fig. 3G; **Table 9**) and Sanger sequencing.

### Protein fractionation and western blot analysis

Parasites were washed once with Dulbecco’s phosphate-buffered saline (DPBS, Thermo Fisher 14190), then resuspended in cytoplasmic lysis buffer (25 mM Tris–HCl pH 7.5, 10 mM NaCl, 1.5 mM MgCl2, 1% IGEPAL CA-630, and 1× protease inhibitor cocktail [“PI”, Roche 4693132001]) at 4°C and incubated on ice for 30 min. The cytoplasmic lysate was cleared with centrifugation (13,500 g, 10 min, 4°C). The pellet (containing the nuclei) was resuspended in 3.3 times less volume of nuclear extraction buffer (25 mM Tris–HCl pH 7.5, 600 mM NaCl, 1.5 mM MgCl2, 1% IGEPAL CA-630, PI) than cytoplasmic lysis buffer at 4°C, transferred to 1.5 mL sonication tubes (Diagenode C30010016, 300 µL per tube), and sonicated for five min total (10 cycles of 30 s on/off) in a Diagenode Pico Bioruptor at 4°C. This nuclear lysate was cleared with centrifugation (13,500 g, 10 min, 4°C). Protein samples were supplemented with NuPage Sample Buffer (Thermo Fisher NP0008) and NuPage Reducing Agent (Thermo Fisher NP0004) and denatured for 10 min at 70°C. Proteins were separated on a 4–12% Bis-Tris NuPage gel (Thermo Fisher NP0321) and transferred to a PVDF membrane with a Trans-Blot Turbo Transfer system (Bio-Rad). The membrane was blocked for 1 h with 1% milk in PBST (PBS, 0.1% Tween 20) at 25°C. HA-tagged proteins, GFP-tagged proteins, and histone H3 were detected with anti-HA (Abcam ab9110, 1:1,000 in 1% milk-PBST), anti-GFP (Chromotek PABG1) and anti-H3 (Abcam ab1791, 1:2,500 in 1% milk-PBST) primary antibodies, respectively, followed by donkey anti-rabbit secondary antibody conjugated to horseradish peroxidase (“HRP”, Sigma GENA934, 1:5,000 in 1% milk-PBST). For some blots, HA-tagged antibodies were detected with an anti-HA-HRP antibody (Cell Signaling C29F4 HRP Conjugate #14031). HRP signal was developed with SuperSignal West Pico Plus or Femto chemiluminescent substrate (Thermo Fisher 34580 or 34096, respectively) and imaged with a ChemiDoc XRS+ (Bio-Rad).

### Chromatin immunoprecipitation and sequencing

Clonal populations of AP2-P-3HA-*glmS* (two replicates: 8 x 10^8^ and 7 x 10^8^) and MORC-3HA-glmS (one replicate: 2.5 x 10^9^) parasites were tightly synchronized and harvested at 36 hpi. Parasite culture was centrifuged at 800 *g* for 3 min at 25°C. Medium was removed and the RBCs were lysed with 10 mL 0.075% saponin (Sigma S7900) in DPBS at 37°C. The parasites were centrifuged at 3,250 *g* for 3 min at 25°C and washed with 10 mL DPBS at 37°C. For the AP2-P-3HA-*glmS* parasites, the supernatant was removed, and the parasite pellet was resuspended in 10 mL of PBS at 25°C. The parasites were cross-linked by adding methanol-free formaldehyde (Thermo Fisher 28908) (final concentration 1%) and incubating with gentle agitation for 10 min at 25°C. The cross-linking reaction was quenched by adding glycine (final concentration 125 mM, Sigma G8899) and incubating with gentle agitation for 5 min at 25°C. For the MORC-3HA-*glmS* parasites, the supernatant was removed, and the parasite pellet was resuspended in 20 mL of PBS at 25°C. MgCl_2_ (Invitrogen AM9530G) was added to a final concentration of 1 mM. 80 μL of ChIP Cross-link Gold (Diagenode C01019027) were added and sample was incubated at 25°C for 30 min with gentle agitation. Parasites were centrifuged at 3,250 *g* for 5 min at 4°C and the supernatant removed. The pellet was washed twice with DPBS at 4°C and centrifuged at 3,250 *g* for 5 min at 4°C. The parasites were resuspended in 20 mL DPBS at 4°C and were further cross-linked by adding methanol-free formaldehyde (Thermo Fisher 28908) (final concentration 1%) and incubating with gentle agitation for 15 min at 25°C. The cross-linking reaction was quenched by adding glycine (final concentration 125 mM, Sigma G8899) and incubating with gentle agitation for 5 min at 25°C. Parasites were centrifuged at 3,250 *g* for 5 min at 4°C and the supernatant removed. Parasite pellets were snap frozen and stored at -80°C.

For each time-point, 200 µL of Protein G Dynabeads (Invitrogen 10004D) were washed twice with 1 mL ChIP dilution buffer (16.7 mM Tris–HCl pH 8, 150 mM NaCl, 1.2 mM EDTA pH 8, 1% Triton X-100, 0.01% SDS) using a DynaMag magnet (Thermo Fisher 12321D). The beads were resuspended in 1 mL ChIP dilution buffer with 8 μg of anti-HA antibody (Abcam ab9110) and incubated on a rotator at 4°C for 6 h.

The cross-linked parasites were resuspended in 4 mL of lysis buffer (10 mM HEPES pH 8, 10 mM KCl, 0.1 mM EDTA pH 8, PI) at 4°C, and 10% Nonidet-P40 was added (final concentration 0.25%). The parasites were lysed in a prechilled dounce homogenizer (100 strokes). The lysates were centrifuged for 10 min at 13,500 *g* at 4°C, the supernatant was removed, and the pellet was resuspended in 3.6 mL SDS lysis buffer (50 mM Tris–HCl pH 8, 10 mM EDTA pH 8, 1% SDS, PI) at 4°C. The liquid was distributed into 1.5 mL sonication tubes (Diagenode C30010016, 300 µL per tube) and sonicated for 12 min total (24 cycles of 30 s on/off) in a Diagenode Pico Bioruptor at 4°C. The sonicated extracts were centrifuged at 13,500 *g* for 10 min at 4°C and the supernatant, corresponding to the chromatin fraction, was kept. The DNA concentration for each time point was determined using the Qubit dsDNA High Sensitivity Assay Kit (Thermo Fisher Scientific Q32851) with a Qubit 3.0 Fluorometer (Thermo Fisher Scientific). For each time point, chromatin lysate corresponding to 100 ng of DNA was diluted in SDS lysis buffer (final volume 200 μL) and kept as “input” at −20°C. Chromatin lysate corresponding to 8 μg for AP2-P-3HA-*glmS* and 10 μg for MORC-3HA of DNA was diluted 1:10 in ChIP dilution buffer at 4°C.

Using a DynaMag magnet, the antibody-conjugated Dynabeads were washed twice with 1 mL ChIP dilution buffer and resuspend in 100 μL of ChIP dilution buffer at 4°C. Then the washed antibody-conjugated Dynabeads were added to the diluted chromatin sample and incubated overnight with rotation at 4°C. The beads were collected on a DynaMag into eight different tubes per sample, the supernatant was removed, and the beads in each tube were washed for 5 min with gentle rotation with 1 mL of the following buffers, sequentially:

`o` Low salt wash buffer (20 mM Tris–HCl pH 8, 150 mM NaCl, 2 mM EDTA pH 8, 1% Triton X-100, 0.1% SDS) at 4°C.
`o` High salt wash buffer (20 mM Tris–HCl pH 8, 500 mM NaCl, 2 mM EDTA pH 8, 1% Triton X-100, 0.1% SDS) at 4°C.
`o` LiCl wash buffer (10 mM Tris–HCl pH 8, 250 mM LiCl, 1 mM EDTA pH 8, 0.5% IGEPAL CA-630, 0.5% sodium deoxycholate) at 4°C.
`o` TE wash buffer (10 mM Tris–HCl pH 8, 1 mM EDTA pH 8) at 25°C.

After the washes, the beads were collected on a DynaMag, the supernatant was removed, and the beads for each time point were resuspended in 800 μL of elution buffer and incubated at 65°C for 30 min with agitation (1000 rpm 30 s on/off). The beads were collected on a DynaMag and the eluate, corresponding to the “ChIP” samples, was transferred to a different tube.

For purification of the DNA, both “ChIP” and “Input” samples were incubated for approximately 10 h at 65°C to reverse the crosslinking. 200 μL of TE buffer followed by 8 μL of RNaseA (Thermo Fisher EN0531) (final concentration of 0.2 mg/mL) were added to each sample, which was then incubated for 2 h at 37 °C. 4 μL Proteinase K (New England Biolabs P8107S) (final concentration of 0.2 mg/mL) were added to each sample, which was then incubated for 2 h at 55°C. 400 μL phenol:chloroform:isoamyl alcohol (25:24:1) (Sigma, 77617) were added to each sample, which was then mixed with vortexing and centrifuged for 10 min at 13,500 *g* at 4°C to separate phases. The aqueous top layer was transferred to another tube and mixed with 30 μg glycogen (Thermo Fisher 10814) and 5M NaCl (200 mM final concentration). 800 μL 100% EtOH at 4°C were added to each sample, which was then incubated at −20°C for 30 min. The DNA was pelleted by centrifugation for 10 min at 13,500 *g* at 4°C, washed with 500 μL 80% EtOH at 4°C, and centrifuged for 5 min at 13,500 *g* at 4°C. After removing the EtOH, the pellet was dried at 25 °C and all DNA for each sample was resuspended in 30 μL 10 mM Tris– HCl, pH 8 total. The DNA concentration and average size of the sonicated fragments was determined using a DNA high sensitivity kit and the Agilent 2100 Bioanalyzer. Libraries for Illumina Next Generation Sequencing were prepared with the MicroPlex library preparation kit (Diagenode C05010014), with KAPA HiFi polymerase (KAPA biosystems) substituted for the PCR amplification. Libraries were sequenced on the NextSeq 500 platform (Illumina).

### RNA extraction and stranded RNA sequencing

An AP2-P-3HA-*glmS* clone was synchronized simultaneously and the culture was split into two at 12 hpi. Glucosamine (Sigma G1514, final concentration 2.5 mM) was added to one culture for two rounds of parasite replication (approximately 96 h). Parasites were then re-synchronized and three technical replicates (with and without glucosamine) were harvested at 36 hpi. RBCs were lysed in 0.075 % saponin (Sigma S7900) in PBS at 37°C, centrifuged at 3,250 *g* for 5 min, washed in PBS, centrifuged at 3,250 g for 5 min, and resuspended in 700 μL QIAzol reagent (Qiagen 79306). RNA was extracted using an miRNeasy Mini kit (Qiagen 1038703) with the recommended on-column DNase treatment. Total RNA was poly (A) selected using the Dynabeads mRNA Purification Kit (Thermo Fischer Scientific 61006). Library preparation was performed with the NEBNext® Ultra™ II Directional RNA Library Prep Kit for Illumina® (New England Biolabs E7760S) and paired end sequencing was performed on the Nextseq 500 platform (Illumina).

### Parasite growth assay

Parasite growth was measured as described previously^89^. Briefly, two AP2-P-3HA-*glms* clones and a WT clone were tightly synchronized. Each culture was split, and glucosamine (Sigma G1514, 2.5 mM final concentration) was added to one half for approximately 96 h before starting the growth curve. The parasites were tightly re-synchronized and diluted to ∼0.2% parasitemia (5% hematocrit) at ring stage. The growth curve was performed in a 96-well plate (200 μL culture per well) with three technical replicates per condition per blood. Every 24 h, 5 μL of the culture were fixed in 45 μL of 0.025% glutaraldehyde in PBS for 1 h at 4°C. After centrifuging at 800 g for 5 min, free aldehyde groups were quenched by re-suspending the iRBC pellet in 200 μL of 15 mM NH4Cl in PBS. A 1:10 dilution of the quenched iRBC suspension was incubated with Sybr Green I (Sigma S9430) to stain the parasite nuclei. Quantification of the iRBCs was performed in a CytoFLEX S cytometer (Beckman Coulter) and analysis with FlowJo Software. **Computational processing and analysis**

### Micro-C processing and analysis

Micro-C paired-end reads (150 bp paired end) were mapped to the *P. falciparum* genome^48^ (plasmoDB.org, version 3, release 55) and processed into pairs and multi-resolution normalized contact matrix .mcool files using hicstuff (https://github.com/koszullab/hicstuff) with the following non-default options: ‵--enzyme mnase --mapping iterative --duplicates --binning 100‵. For each condition, correlation between replicates was estimated using hicrep^90^.

Replicates showed good correlation; their pairs files were subsequently merged using pairtools^91^ and multi-resolution binned contact matrix files .mcool files of merged replicates were regenerated using cooler^92^. When comparing WT to AP2-P KD Micro-C results, only WT replicates generated with the same genetic background as the AP2-P KD were used, and WT contacts were subsampled with pairtools to 65.106 contacts to match AP2-P KD Micro-C data. Hi-C data was imported and manipulated in R using HiCExperiment^93^. Genomic distance-dependent contact frequency was computed using HiContacts^93^. O/E contact frequency was computed using the ‵detrend()‵ function from HiContacts^93^. Aggregated contact maps were generated using the ‵aggregate()‵ function from HiContacts^93^. All contact maps, including aggregate plots, were generated in R using HiContacts^93^.

Insulation score for any given genomic locus of interest (e.g., a HP1 domain border) was estimated by dividing the summed frequency of contacts within a 20kb window centered at the genomic locus of interest that are not spanning this locus, by the total summed frequency of contacts within this 20kb window. Higher scores indicate higher insulation of the genomic locus of interest.

Boundaries and loops were annotated in WT Micro-C data using the automated structural feature caller chromosight^94^, using 2 kb resolution for boundaries and 1 kb resolution for loops and filtering for q-values ≤ 10^-4^. Boundaries were split into “euchromatin” (211 boundaries), “HP1 domain”, “Virulence-coding subtelomeres” or “non-genic subtelomeres” according to their genomic location (Table 6), and their insulation score was computed as described above. Loops were filtered to only retain those anchored at identified euchromatic boundaries (Table 7). 209 protein-coding genes were found overlapping a euchromatin boundary, and 97 were found overlapping a euchromatic loop anchor.

An overlap between a euchromatic protein-coding gene and AP2-P, AP2-I and/or MORC peaks was considered if peaks were ≤ 1kb away from that gene. The enrichment in AP2-P, AP2-I and/or MORC peaks over euchromatic protein-coding genes was compared to the rest of protein-coding genes using Fisher’s exact test.

### Hi-C processing and analysis

Two publicly available Hi-C datasets^27,66^ were reprocessed using hicstuff with the following options: ‵--enzyme MboI --mapping iterative --duplicates --binning 100‵.

### ChIP-seq processing and analysis

Sequenced reads (150 bp paired end) were mapped to the *P. falciparum* genome^48^ (plasmoDB.org, version 3, release 56) using bowtie^95^. PCR duplicates were filtered using samtools’ fixmate and markdup^96^ commands and only alignments with a mapping quality ≥ 30 were retained (samtools view -q 30)^96^. The paired end deduplicated ChIP and input BAM files were used as treatment and control, respectively, for peak calling with the MACS2 subcommands^97^. In brief, for each ChIP experiment, pileup files were first generated using the MACS2 pileup command, and the larger of the two files (i.e., Input or control) was down-sampled using MACS2 bdgopt. q-values and fold enrichment of ChIP/input was then calculated using MACS2 bdgcmp. Final peak calling was performed using MACS2 bdgcallpeak using a q-value cut-off of 0.001 (-c 3). For the two biological replicates of AP2-P, consensus peaks shared between biological replicates 1 and 2 were defined using the bedtools intersect command^98^. ChIP/input ratio tracks were generated using deeptool’s bamCompare command^99^. Integrative Genomics Viewer^100^ was used to inspect tracks and MACS2 peaks.

Binding peaks were associated to the nearest protein-coding genes using bedtools closest command^98^ along with *P. falciparum* reference genome feature file (gff) (plasmoDB.org, version 3, release 56). Only regions 500 bp upstream or downstream near to the protein coding genes were considered further for downstream analysis. To perform functional analysis on the genes closest to the peaks, Gene Ontology Enrichment tool from PlasmoDB web interface (plasmoDB.org, version 3, release 56) was used for Ontology term – Biological Process with a *P*-Value cutoff of 0.05. Fold change quantification and statistical analysis for all peaks and peaks in centromeric regions was performed in R^101^.

Metagene plots in **Figs. 2C, 4B**, and **Supp.** Fig. 4B were generated using the plotAvgProf2 command in the ChIPSeeker package^102^. ChIP/Input ratios were calculated genome-wide using ‘bamCompare’ and normalized across regions of interest using ‘computeMatrix’ in the deeptools package^99^. Final matrices were plotted using ‘plotProfile’. The tidyCoverage package was also used to generate coverage heatmaps and aggregate coverage tracks in R^103^.

*de novo* motif discovery analysis was performed using MEME-ChIP suite^104^. ChIP-seq peak summits identified using MACS2 were extended +/-100 bp and used as the input. These extended summits were then converted to fasta format using bedtools getfasta command^98^. The resulting file was used as an input for the MEME-ChIP motif search algorithm.

Pie charts were made in R^101^. Venn diagram was made with DeepVenn^105^.

### RNA-seq processing and analysis

Sequenced reads were mapped to the *P. falciparum* genome (plasmoDB.org, version 3, release 56) using “STAR”^106^, restricting the number of multiple alignments allowed for a read with (option --outFilterMultimapNmax 1”). Alignments were subsequently filtered for duplicates and a mapping quality ≥ 20 using samtools^96^. Gene counts were quantified with htseq-count^107^ and differentially expressed genes were identified in R using package DESeq2^108^. Gene Ontology enrichment analysis was performed on differentially expressed genes (*q* < 0.05) using the built-in tool at PlasmoDB.org^109^ (version 3, release 56) with default settings for Biological Process (*P*-value < 0.05).

RNA-seq-based cell cycle progression was estimated in R^101^ by comparing the normalized expression values (i.e., RPKM, reads per kilobase per exon per one million mapped reads) of each sample to the microarray data from^25^ using the statistical model as in^28^.

Barplots and dotplots were made using ggplot package in R^101^. Histograms were made using the data from^63^.

## Data Accessibility

All data sets (ChIP-seq, RNA-seq, Micro-C) generated in this study will be available upon peer-reviewed publication.

Previously published data sets utilized in this study are available at the following NCBI accession numbers:

- AP2-I ChIP from (Santos *et al*., 2017)^62^: SRR5114665
- AP2-I ChIP Input from (Santos *et al*., 2017)^62^: SRR5114667
- TRZ ChIP from (Bertschi *et al*., 2017)^60^: SRR3085676
- TRZ ChIP Input from (Bertschi *et al*., 2017)^60^: SRR3085677
- HP1 ChIP from (Carrington *et al*., 2021)^110^: SRR12281320
- HP1 ChIP Input from (Carrington *et al*., 2021)^110^: SRR12281322
- ATAC-seq from (Toenhake *et al*., 2018)^58^: SRR6055333
- ATAC-seq gDNA control from (Toenhake *et al*., 2018)^58^: SRR6055335
- Hi-C from (Ay *et al*., 2014)^27^: SRR957166
- Hi-C from (Subudhi *et al*., 2023)^66^: SRR19611536

## Figure Legends

**Supplementary Figure 1:**
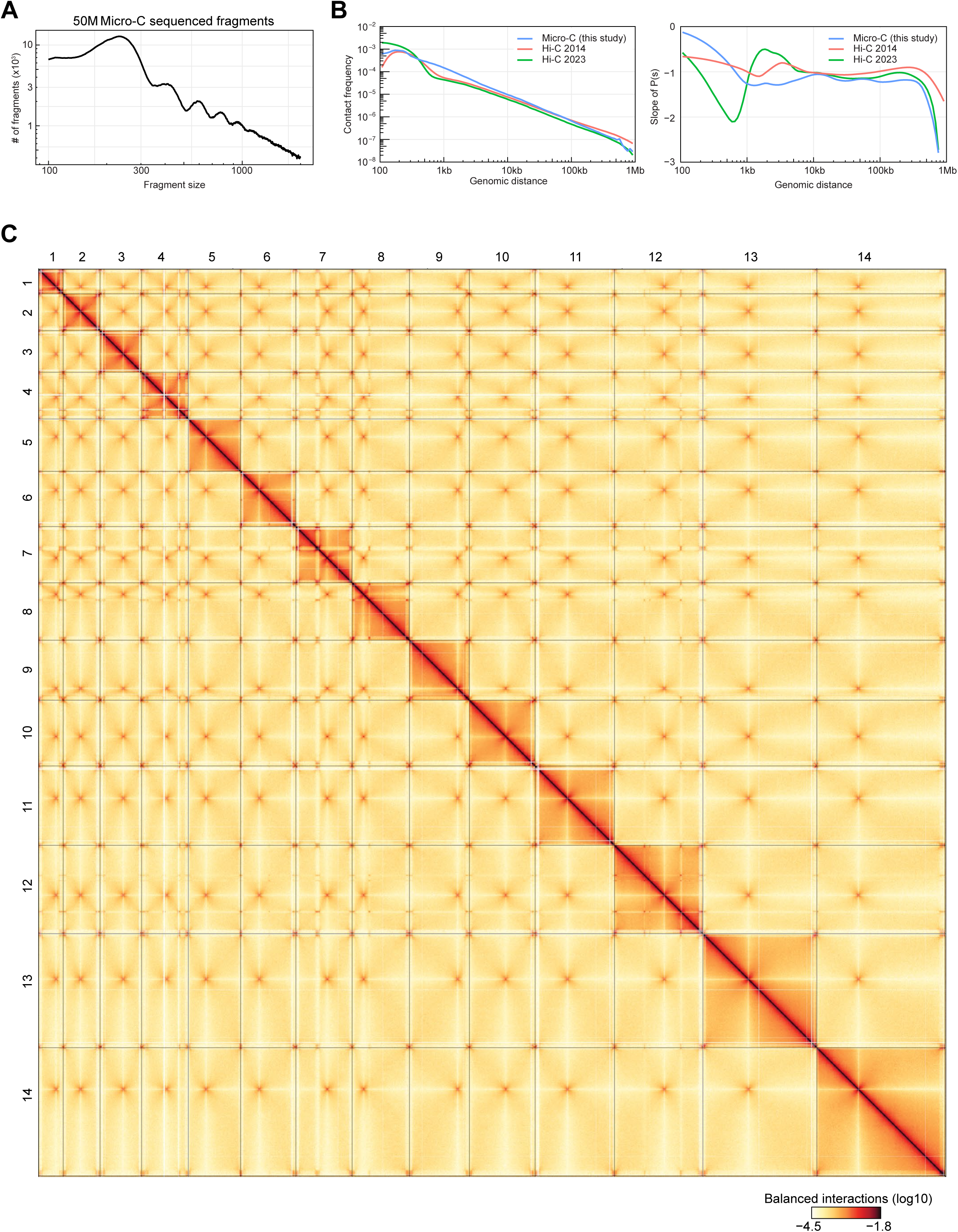
Quality control of the Micro-C technique in *P. falciparum*. **A)** Fragment size distribution in the Micro-C library. Only 50 x 10^6^ fragments have been used. **B)** Comparison of genomic distance-dependent interaction frequencies in Micro-C data (this study) and previously published Hi-C datasets^27,66^ from *P. falciparum*. Left: P(s); right: slope of P(s). **C)** Micro-C contact map across all *P. falciparum* chromosomes.

**Supplementary Figure 2:**
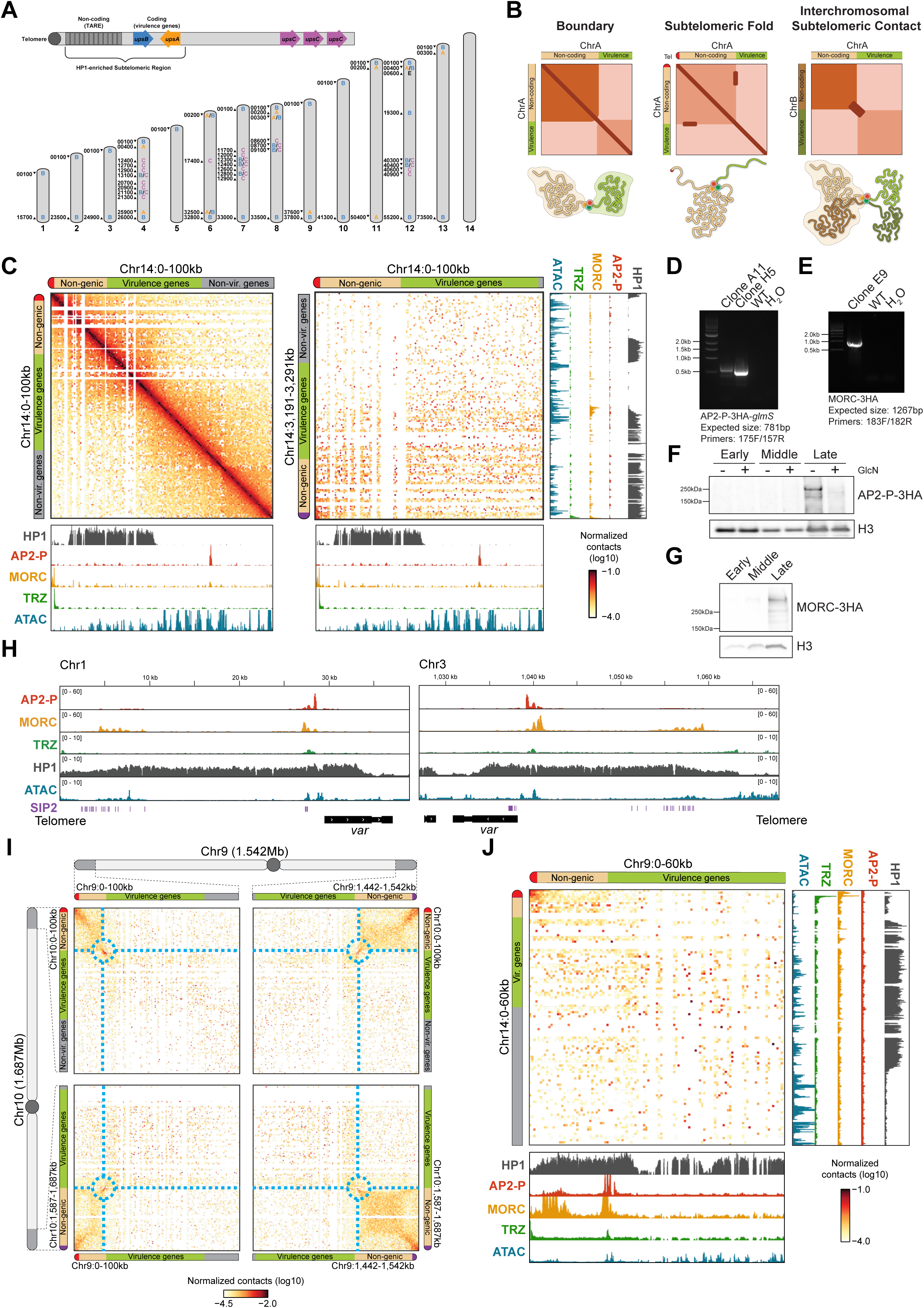
Micro-C reveals subtelomeric fold structures defined by a multiprotein complex. **A)** Schematic of how *var* genes are categorized and located within a chromosome (top) and throughout the genome (bottom). Subtelomeric *var* genes are indicated in blue (*upsB*) and orange (*upsA*), and central *var* genes are indicated in purple (*upsC*). Chromosomes are represented with gray bars, and chromosome number is indicated under each bar. *var* gene type is indicated on the chromosome, and gene ID (excluding the preceding chromosome number) is listed to the left of its position on the chromosome. The direction of *var* gene transcription is indicated with an arrowhead. **B)** Graphical representation of the different structures observed with Micro-C in subtelomeric regions. Top: Theoretical contact maps, with darker colors indicating higher frequency of contacts. Red is the left arm telomere, tan is the non-genic subtelomeric region, and green is the virulence gene-encoding subtelomeric region. Bottom: Illustrations of hypothetical chromosomal conformations corresponding to the contact maps above. Red balls are AP2-P, yellow balls are MORC, and green balls are TRZ. **C)** Micro-C contact maps of the end of Chromosome 14 left arm (left) or between the telomere of each arm of chromosome 14 (right) in late-stage parasites (100kb-wide, 1kb resolution). AP2-P, MORC, HP1^110^, and TRZ^60^ ChIP/input ratio tracks as well as ATAC-seq^58^ track from late-stage parasites are shown at the bottom and to the right of contact maps. To the top and left of each contact map: red is the left arm telomere, tan is the non-genic subtelomeric region, green is the virulence gene-encoding subtelomeric region, grey is non-virulence genes, and purple is the right arm telomere. **D)** and **(E)** DNA gels showing PCR validation of the AP2-P-3HA-*glmS* (D) and MORC-3HA (E) strains with the indicated primers (**Table 9**). Bands indicate integration of the 3HA-*glmS*-encoding and 3HA-encoding sequences at the 3’ ends of the endogenous *ap2-p* or *morc* genes, respectively, in the clones used in this study. No genomic DNA (H_2_O) and genomic DNA from WT parasites (WT) were used as controls. DNA size is indicated with a ladder at the left side of each gel, and expected band sizes are indicated at the bottom of each gel. **(F)** Western blot analysis of nuclear extracts from a synchronous clonal population of AP2-P-3HA-*glmS* parasites at early, middle, and late stages in the absence (-) or presence (+) of glucosamine (GlcN). Parasites were treated with glucosamine for 48 hr. AP2-P-3HA was detected with an anti-HA antibody, and an antibody against histone H3 was used as a loading control. Molecular weights are shown to the left. **(G)** Western blot analysis of nuclear extracts from a synchronous clonal population of MORC-3HA parasites at early, middle, and late stages. MORC-3HA was detected with an anti-HA antibody, and an antibody against histone H3 was used as a loading control. Molecular weights are shown to the left. **(H)** ChIP-seq data showing enrichment (*y*-axis = ChIP/Input) of AP2-P, MORC, TRZ^60^, and HP1^110^ in late-stage parasites at the subtelomeric region on the left arm of chromosome 1 and the right arm of chromosome 3. ATAC-seq data^58^ from a closely corresponding stage (35 hpi) shows chromatin accessibility [*y*-axis = ATAC-seq (RPM)/gDNA (RPM)]. SIP2 binding sites (SPE2 sequence)^59^ are indicated with vertical purple lines. The *x*-axis is DNA sequence, with the *var* gene represented by a black box with white arrowheads to indicate transcription direction. The telomere is also indicated. **(I)** Interchromosomal Micro-C contact map between subtelomeric regions of either arm of chromosome 10 and either arm of chromosome 9 in late-stage parasites (100kb-wide, 1kb resolution). **(J)** Interchromosomal Micro-C contact map between subtelomeric region of the left arm of chromosome 9 and the left arm of chromosome 14 in late-stage parasites (60kb-wide, 1kb resolution). AP2-P, MORC, HP1^110^, and TRZ^60^ ChIP/input ratio tracks as well as ATAC-seq^58^ track from late-stage parasites are shown at the bottom and to the right of contact maps.

**Supplementary Figure 3:**
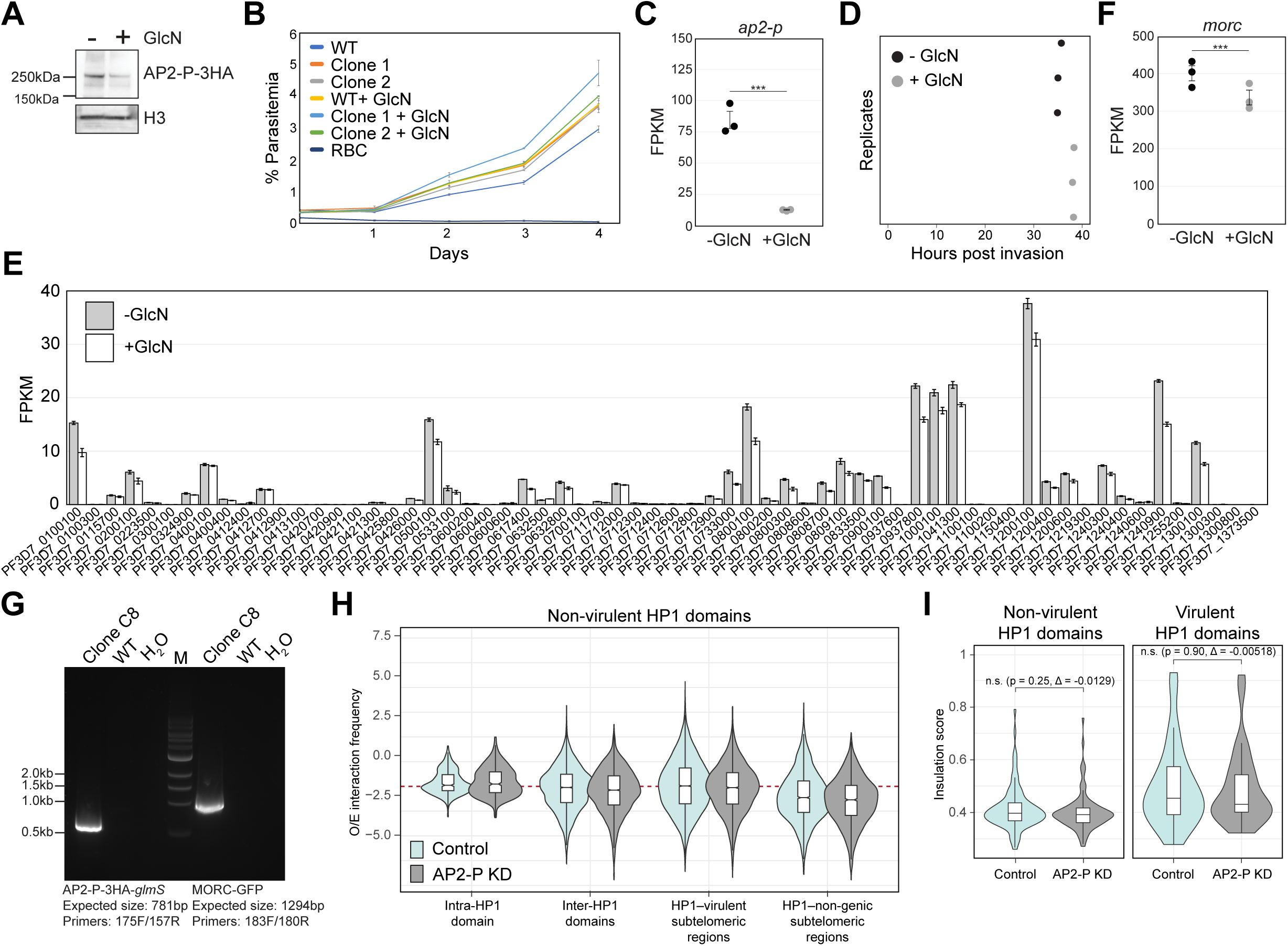
AP2-P KD leads to MORC KD, but no phenotype in growth, *var* gene transcription, interactions amongst non-virulent HP1 domains, or insulation of HP1 domains. **(A)** Western blot analysis of nuclear extracts from a synchronous clonal population of AP2-P-3HA-*glmS* late-stage parasites in the absence (-) or presence (+) of glucosamine (GlcN). Parasites were treated with glucosamine for 48 hr. AP2-P-3HA was detected with an anti-HA antibody, and an antibody against histone H3 was used as a loading control. Molecular weights are shown to the left. **(B)** Growth curve showing parasitemia over four days for WT and two clones of AP2-P-3HA-*glmS* parasites in the absence or presence of glucosamine (GlcN). Glucosamine treatment was started two cycles (96 h) before Day 0 to ensure AP2-P KD during the days sampled. Uninfected red blood cells (RBC) served as reference of background. Error bars indicate standard deviation of three technical replicates (n = 3). A two-way ANOVA with Tukey post hoc test was used for statistical analysis. No significant differences were found. **(C)** RNA-seq of an AP2-P-3HA-*glmS* clone shows *ap2-p* transcript levels (FPKM) in late-stage parasites (*q* = 4 x 10^-4^) in the absence (black circles) or presence (grey circles) of glucosamine (GlcN). Error bars represent standard deviation of three technical replicates. P-values are calculated with a Wald test for significance of coefficients in a negative binomial generalized linear model as implemented in DESeq2^108^. *q* = Bonferroni corrected P-value. Asterisks indicate a *q* value < 0.05. Corresponding data can be found in (**Table 2**). **(D)** Cell cycle progression (hours post invasion on x-axis) estimation of synchronous, clonal AP2-P-3HA-*glmS* populations in the absence or presence of glucosamine (GlcN). Parasites harvested at 36 hpi were compared to microarray data from^25^ as in^111^ to determine the approximate time point in the red blood cell cycle (x-axis). Replicates are represented with black (−GlcN) or grey (+GlcN) circles. **(E)** RNA-seq of an AP2-P-3HA-*glmS* clone shows all *var* gene transcript levels (FPKM) in late-stage parasites in the absence (grey) or presence (white) of glucosamine (GlcN). Error bars represent standard deviation of three technical replicates. P-values are calculated with a Wald test for significance of coefficients in a negative binomial generalized linear model as implemented in DESeq2^108^. *q* = Bonferroni corrected P-value. Corresponding data can be found in (**Table 2**). **(F)** RNA-seq of an AP2-P-3HA-*glmS* clone shows *morc* transcript levels (FPKM) in late-stage parasites (*q* = 0.0035) in the absence (black circles) or presence (grey circles) of glucosamine (GlcN). Error bars represent standard deviation of three technical replicates. P-values are calculated with a Wald test for significance of coefficients in a negative binomial generalized linear model as implemented in DESeq2^108^. *q* = Bonferroni corrected P-value. Asterisks indicate a *q* value < 0.05. Corresponding data can be found in (**Table 2**). **(G)** DNA gel showing PCR validation of the AP2-P-3HA-*glmS*:MORC-GFP strain with the indicated primers (**Table 9**). Bands indicate integration of the 3HA-*glmS*-encoding and GFP-encoding sequences at the 3’ ends of the endogenous *ap2-p* or *morc* genes, respectively, in the clones used in this study. No genomic DNA (H_2_O) and genomic DNA from WT parasites (WT) were used as controls. DNA size is indicated with a ladder at the left side of the gel, and expected band sizes are indicated at the bottom. **(H)** Log_2_-scaled observed/expected interaction frequency ratio for (i) intra-chromosomal interactions between two non-virulence gene-containing HP1 domains, (ii) interchromosomal interactions between two non-virulence gene-containing HP1 domains, (iii) interactions between non-virulence gene-containing HP1 domains and virulence gene-coding subtelomeric loci, or (iv) interactions between non-virulence gene-containing HP1 domains and non-genic subtelomeric loci. Contacts in the subsampled control dataset are shown in cyan and those in the AP2-P KD dataset are shown in grey. **(I)** Insulation score at borders of non-subtelomeric HP1 domains, split according to whether each domain contains virulence genes. Contacts in the subsampled control dataset are shown in cyan and those in the AP2-P KD dataset are shown in grey.

**Supplementary Figure 4:**
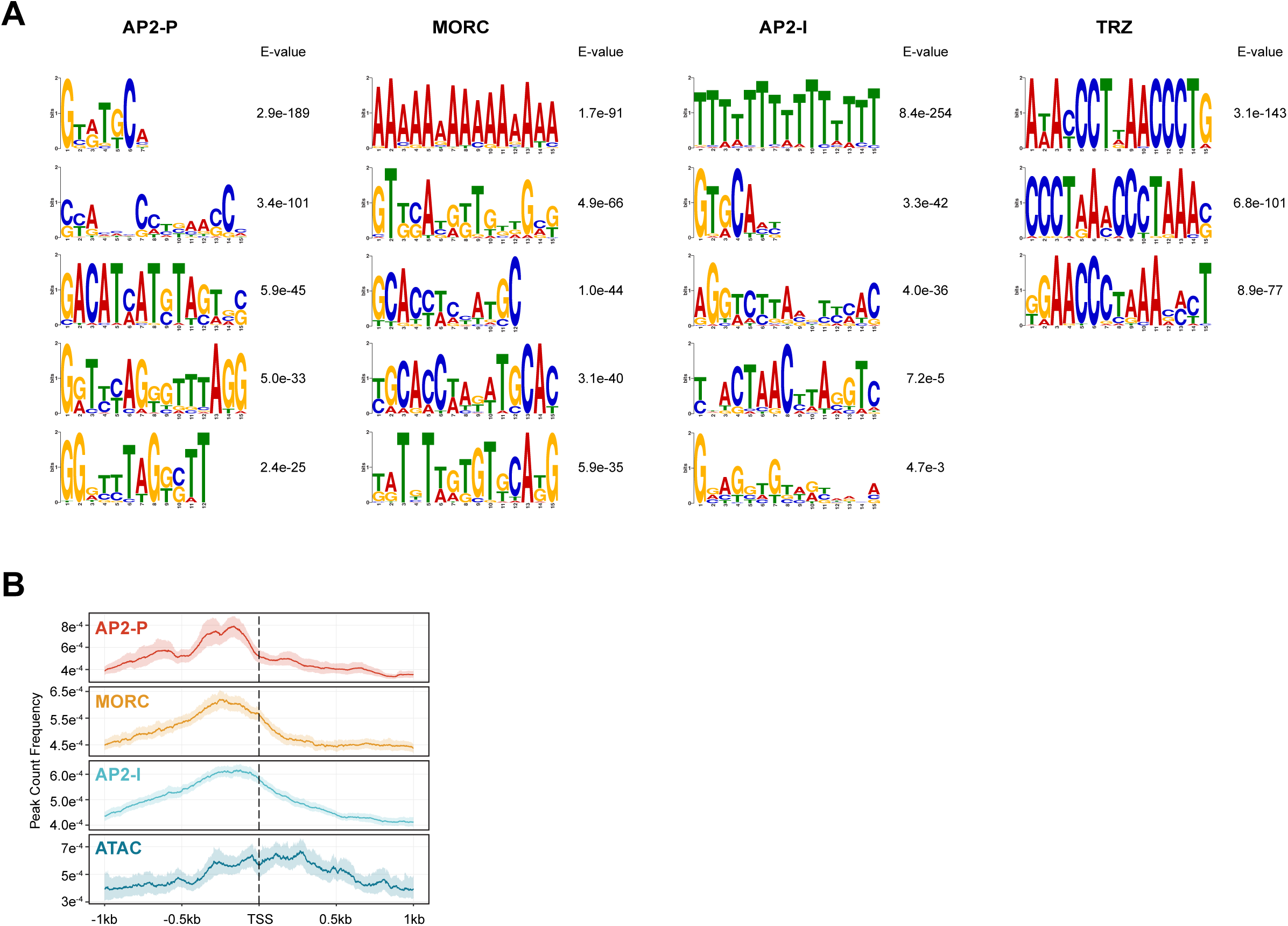
AP2-P, MORC, AP2-I, and TRZ binding patterns. **A)** Motif discovery analysis of AP2-P, MORC, AP2-I^62^, and TRZ^60^ ChIP-seq peaks. Top hits with minimum threshold E-value < 0.05 are shown. **B)** Metagene plots showing average AP2-P and MORC enrichment (y-axis = peak count frequency) in clonal AP2-P-3HA-*glmS* and MORC-3HA parasites, respectively, in late-stage parasites from 1 kb upstream of the TSS to 1 kb downstream of all genes that are significant downregulated more than two-fold upon AP2-P KD. AP2-I ChIP-seq data^62^ and ATAC-seq data^58^ from stage-matched parasites is included. One replicate was used for the AP2-P and MORC ChIP datasets.

**Supplementary Figure 5:**
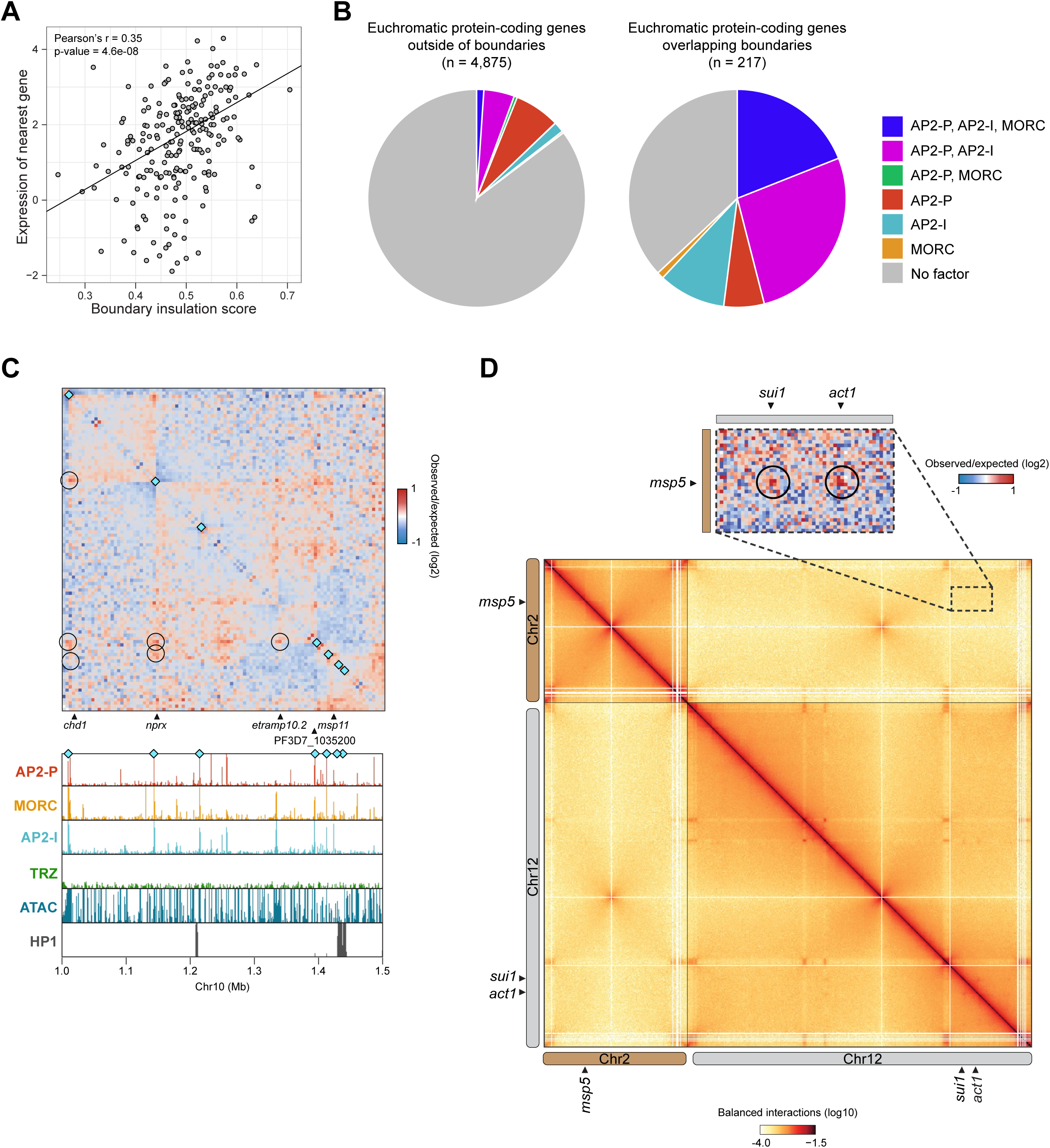
AP2-P activating complex defines euchromatic boundaries and long-range interactions. **(A)** Pearson correlation between insulation score of 211 euchromatic boundaries and the level of expression of the nearest gene. **(B)** Pie charts showing the proportion of euchromatic protein-coding genes outside (left) and overlapping (right) Micro-C boundaries that are associated with a peak of AP2-P, MORC, and/or AP2-I (peak ≤ 1kb away from gene). **(C)** Micro-C contact map of a central region of chromosome 10 in late-stage parasites. Micro-C boundaries are indicated with cyan diamonds. Selected genes are indicated at the bottom. AP2-P and MORC ChIP/Input ratio signals from late-stage parasites is shown at the bottom. Stage-matched AP2-I^62^, TRZ^60^, HP1^110^ ChIP-seq and ATAC-seq^58^ data are included. Long-range loops identified with chromosight^94^ are indicated with black circles on the contact map. **(D)** Micro-C contact maps focusing on intra- and interchromosomal contacts in chromosomes 2 and 12. Interchromosomal interactions between the *msp2* locus (Chr2) and *sui1* or *act1* loci (Chr12) are highlighted in the inset showing observed/expected contact frequency ratio.

## Table Legends

**Table 1:** Peaks of AP2-P (consensus peaks of two biological replicates), MORC, AP2-I^62^, or TRZ^60^ in late-stage parasites and their chromosomal coordinates, significance [-log_10_(*q*-value)], summit coordinate, fold enrichment (FE) at the summit, the closest unique protein-coding gene, the peak’s relative position to the closest gene (upstream, coding sequence, 3’ UTR, 5’ UTR, or downstream), and the distance to the closest gene feature.

**Table 2:** Differential gene expression analysis in late-stage parasites (36 hpi) of glucosamine-treated over untreated AP2-P-3HA-*glmS* parasites. Shown is the gene ID, baseMean, log_2_(FoldChange), *P* value, and adjusted P value. Genes are sorted in ascending order of adjusted *P* value. PF3D7_1107800, which encodes AP2-P, is highlighted in grey.

**Table 3:** Gene Ontology analysis (biological process) for genes with a peak of AP2-P in their upstream or 5’UTR region in late-stage parasites (defined in Table 1). Shown are Gene Ontology ID, name of category, total number of genes in this category (Bgd count), number of genes from the query in this category (Result count), genes from the query in this category (Result gene list), percentage of genes from the query in this category out of total number of genes in this category (Pct of bgd), *P* value, and Bonferroni adjusted *P* value.

**Table 4:** Genes whose products play a role in parasite egress from or invasion into the red blood cell^64,65^.

**Table 5:** Gene Ontology analysis (biological process) for genes that are significantly downregulated more than two-fold upon AP2-P KD in late-stage parasites (defined in Table 2). Shown are Gene Ontology ID, name of category, total number of genes in this category (Bgd count), number of genes from the query in this category (Result count), genes from the query in this category (Result gene list), percentage of genes from the query in this category out of total number of genes in this category (Pct of bgd), *P* value, and Bonferroni adjusted *P* value.

**Table 6:** Micro-C boundaries in late-stage AP2-P-3HA-*glmS* parasites and their position on each chromosome, *q*-value, insulation score in the absence or presence of glucosamine (GlcN), type of location (non-genic subtelomeric, euchromatic, non-subtelomeric HP1 domain, subtelomeric virulence genes, or telomeric), and the nearest gene (and its product) to the boundary.

**Table 7:** Micro-C long-range contacts between euchromatic genes. Chromosome location, width of, and nearest gene ID are shown for each anchor, as well as the loop interaction score and its P value and *q*-value.

**Table 8:** Gene Ontology analysis (biological process) for euchromatic genes that form long-range interactions (defined in Table 7). Shown are Gene Ontology ID, name of category, total number of genes in this category (Bgd count), number of genes from the query in this category (Result count), genes from the query in this category (Result gene list), percentage of genes from the query in this category out of total number of genes in this category (Pct of bgd), *P* value, and Bonferroni adjusted *P* value.

**Table 9:** Primers, oligos, and DNA blocks used in this study.

**Table 10:** Chromosomal coordinates of subtelomeric fold structures (used to make **Figs. 2B and 3C**).

## Notes

### Competing Interest Statement

The authors have declared no competing interest.

